# Prime Editing Corrects the *HBB* Codon 8/9 (+G) Mutation in Patient-Derived Induced Pluripotent Stem Cells and Restores β-Globin Expression in iPSC-Derived Erythroid Cells

**DOI:** 10.64898/2026.09.22.753453

**Authors:** Irfan Hussain, Kainaat Mumtaz, Maliha Javed, Susheel Fatima, Hijab Zahra, Anwar Alam, Muhammad Jameel, Naveed Altaf Malik, Farhatullah Syed, Fawad Ur Rehman, Salman Kirmani, Ambrin Fatima, Afsar Ali Mian

## Abstract

**Background:** Homozygosity for the *HBB* codon 8/9 (+G) frameshift (c.27dup (p.Ser10ValfsTer14)) causes transfusion-dependent β⁰-thalassaemia and is common in South Asia. Prime editing can reverse this insertion without double-strand breaks or donor DNA, but its efficiency depends on pegRNA design.

**Methods:** We derived Sendai-reprogrammed iPSCs from a homozygous patient, optimised PEmax editing (spacer, pegRNA extension, secondary nick, MLH1dn) and compared patient, PEmax-treated and control iPSC-derived erythroid cells by flow cytometry, colony assays, RT-qPCR, western blotting and cation-exchange HPLC.

**Results:** Patient iPSCs had a normal 46,XY karyotype, expressed pluripotency markers, formed all three germ layers and were Sendai-free by passage 15. A PAM-disrupting spacer with a +68 nicking sgRNA gave the highest intended-edit frequency 8.4%; MLH1dn added little. PEmax-treated cultures matured and formed colonies more like control than patient cultures, restored *HBB* transcript (about 4-fold control iPSC-derived cells) and detectable β-globin protein, and contained an HPLC fraction consistent with HbA (undetectable in patient cells). Fetal haemoglobin (about 80%) and embryonic globin remained predominant.

**Conclusions:** Prime editing corrects *HBB* c.27dupG in patient iPSCs and restores β-globin expression, with partial restoration of adult haemoglobin in a fetal/embryonic-type erythroid background. Validation in additional donors, haematopoietic stem cells and in vivo is required.

## 1. Introduction

Thalassaemia syndromes are recessively inherited haemoglobinopathies in which reduced or absent synthesis of haemoglobin chains results in chronic anaemia of varying severity (Taher et al., 2021). Among the many HBB variants that cause β-thalassaemia, the codon 8/9 (+G) frameshift (*HBB*:c.27dup (p.Ser10ValfsTer14)) is one of the most frequent alleles in Pakistan, accounting for about 16% of alleles in a large Karachi cohort (Ansari et al., 2011), and typically causes a severe, transfusion-dependent β⁰ phenotype from early childhood (Weatherall, 2001; Taher et al., 2021). Standard care comprises regular transfusions and iron chelation, which relieve symptoms without addressing the genetic defect and carry cumulative risks of iron overload, cardiac disease, and endocrinopathy (Taher et al., 2021; Miri-Aliabad, 2025). Allogeneic haematopoietic stem cell transplantation is curative but constrained by donor availability, graft-versus-host disease, and procedure-related morbidity (Angelucci et al., 2014; Anurathapan et al., 2022). Autologous approaches based on gene addition or genome editing of patient-derived cells have therefore emerged as a major therapeutic direction (Bank et al., 2005; Negre et al., 2015; Goodman & Malik, 2016; Niu et al., 2016; Ferrari et al., 2021; Locatelli et al., 2024).

Induced pluripotent stem cell (iPSC) technology enables somatic cells to be reprogrammed into a pluripotent state by ectopic expression of defined transcription factors, generating patient-specific lines that retain the donor genotype (Takahashi & Yamanaka, 2006; Takahashi et al., 2007). Integration-free delivery, for example with Sendai virus or episomal plasmids, avoids genomic integration of the reprogramming factors (Dowey et al., 2012; Beers et al., 2015), and derivation under feeder-free, xeno-free conditions has been described (Chou et al., 2015). Such lines can be genetically corrected, expanded as clonal populations with full genomic quality control, and differentiated into disease-relevant lineages, including haematopoietic progenitors and erythroid cells (Xie et al., 2014; Xu et al., 2015; Ou et al., 2016; Liu et al., 2017). This combination of clonal isolation and unlimited expansion is a distinctive advantage of the iPSC platform for evaluating editing strategies because it allows every candidate genotype to be characterised in depth before differentiation (Ou et al., 2016).

The regulatory approval of exagamglogene autotemcel (exa-cel) in 2023–2024 established the clinical viability of CRISPR-based therapy for β-haemoglobinopathies (Frangoul et al., 2021; Locatelli et al., 2024). That therapy acts by disrupting the erythroid enhancer of BCL11A to induce fetal haemoglobin and does not restore endogenous β-globin synthesis. Approaches that directly correct the causal HBB lesion therefore remain a complementary objective, particularly when the intended mechanism is restoration of the α/β chain balance (Magrin et al., 2019).

Prime editing enables targeted insertions, deletions, and base substitutions without double strand breaks or exogenous donor DNA (Anzalone et al., 2019). The PEmax architecture, which incorporates an optimised Cas9-H840A nickase–reverse transcriptase fusion with improved nuclear localisation, substantially increases editing efficiency across cell types, and transient suppression of mismatch repair by co-expression of dominant-negative MLH1 (MLH1dn; the PE4/PE5 systems) further increases the recovery of intended edits by limiting their reversal (Chen et al., 2021). Prime editing efficiency is nonetheless highly dependent on prime editing guide RNA (pegRNA) architecture, particularly the length of the reverse-transcriptase template (RTT) and primer binding site (PBS) and the position of any secondary nick. These parameters require empirical optimisation at each locus (Doman et al., 2022; Kim et al., 2021; Nelson et al., 2022). *Ex vivo* prime editing of patient-derived haematopoietic stem and progenitor cells has been shown to rescue sickle cell disease phenotypes after engraftment in mice, providing proof of concept for this editing modality in haemoglobinopathies (Everette et al., 2023), and prime editing has been used to install β-thalassaemia mutations in *HBB* in erythroid progenitor cells (Zhang et al., 2022).

Here, we derived iPSCs from dermal fibroblasts of a transfusion-dependent patient homozygous for *HBB* codon 8/9 (+G), characterised them using pluripotency criteria, and systematically optimised PEmax prime editing at this locus across RTT length, secondary nick position, and mismatch repair modulation. We then isolated biallelically corrected clones, assessed predicted off-target sites by Sanger sequencing followed by deep amplicon sequencing, and differentiated corrected and uncorrected lines along the erythroid lineage to compare globin transcript, protein, and haemoglobin tetramer output. To our knowledge, prime editing of this variant has not previously been reported in patient-derived iPSCs with an erythroid functional readout.

## 2. Materials and Methods

### 2.1 Ethical compliance and cell line deposition

The study was approved by the Aga Khan University Ethics Review Committee (2022-6532-22590). Written informed consent was obtained from the β-thalassaemia patient (homozygous *HBB* codon 8/9 (+G)) and from a healthy control donor. All procedures conformed to the Declaration of Helsinki. Established lines are stored at the Centre for Regenerative Medicine and Stem Cell Research at Aga Khan University in accordance with international biobanking standards.

### 2.2 Isolation and culture of patient-derived fibroblasts

Dermal fibroblasts were isolated from a 6-mm skin punch biopsy. The biopsy was cut into approximately 1-mm fragments and placed in a 6-well plate for explant culture. Explants were allowed to adhere for 30–60 min at 37 °C before Dulbecco’s Modified Eagle’s Medium (DMEM) supplemented with 10% fetal bovine serum (FBS; Gibco) and 1% penicillin–streptomycin (Gibco) was added. Cultures were maintained at 37 °C and 5% CO₂, with medium exchanged every 3 days.

Outgrowing fibroblasts were collected after approximately two weeks and expanded to passage three before reprogramming.

### 2.3 Reprogramming of fibroblasts into iPSCs

Fibroblasts were generated and cultured in DMEM (Gibco) supplemented with 10% fetal bovine serum, 1% Penicillin/Streptomycin (Gibco) and 1% Non-essential Amino Acid (NEAA, Gibco) . Fibroblasts of passage 4 were reprogrammed into hiPSCs using Cytotune™-iPS 2.0 Sendai Reprogramming kit (ThermoFisher Scientific) following manufacturer’s guideline. On day 0 of transduction, the cells were transduced using the CytoTune™ 2.0 Sendai reprogramming kit with 4 reprogramming factors at a recommended MOI. After which, the medium was changed every 24 h to remove the virus, and the medium was changed every two days. On day 7 of post-transduction, the cells were transferred to a Matrigel (Thermo Fisher Scientific) coated 6-well plate at a recommended density. After 24 h, the medium was changed to Essential 8™ Medium (Gibco). Two to three weeks after transduction, the appropriate size colonies were manually selected and cultured as iPSCs.

### 2.4 Characterisation of iPSC lines

Candidate colonies were stained with the AP Blue Membrane Substrate Kit (Sigma-Aldrich); AP-positive clones were expanded. Cells were fixed in 4% paraformaldehyde, permeabilised with 0.1% Triton X-100 (omitted for the cell-surface marker TRA-1-60), blocked in 5% BSA, and incubated overnight at 4 °C with primary antibodies against OCT4, SOX2, NANOG, and TRA-1-60, followed by fluorophore-conjugated secondary antibodies and DAPI counterstaining. Images were acquired on a Nikon A1R confocal microscope with identical laser power, gain, and exposure settings across all channels and specimens within an experiment. Single-channel images and merged overlays are presented to allow independent assessment of subcellular localization for each marker.

#### Quantitative RT-PCR for pluripotency markers

Total RNA was extracted using the RNeasy Mini Kit (Qiagen), DNase I-treated, and reverse transcribed with SuperScript™ Mix (Thermo Fisher). Reactions were run with LightCycler® 480 SYBR Green I Master (Roche) for six endogenous pluripotency genes (OCT4, SOX2, NANOG, LIN28, REX1, DNMT3B) in parental fibroblasts, patient iPSCs, and human ESCs, with three technical replicates from three independent RNA preparations. GAPDH served as the reference gene. Data are reported as 2−ΔCt relative to GAPDH and are plotted as a percentage of the ESC value for each gene (Figure 3b).

#### Sendai virus clearance

Residual Sendai vector was quantified by TaqMan® RT-PCR using the TaqMan® iPSC Sendai Detection Kit (A13640; Thermo Fisher), which includes probe assays for the Sendai backbone (SeV), the polycistronic KOS cassette, and the KLF4 and c-MYC transgenes, along with an endogenous control assay for normalisation. Reactions were run on a QuantStudio™ system according to the manufacturer’s protocol (Thermo Fisher MAN0015825). Samples comprised early-passage transduced fibroblasts (positive control), patient iPSCs at passages 2 and 15, and human ESCs (negative control). A sample was classified as vector-free when no amplification was detected within 40 cycles for all four assays, while the endogenous control amplified normally; wells without amplification were assigned Ct = 40 (undetermined) for graphing purposes.

#### Karyotype analysis

G-banding was performed on ≥20 metaphase spreads per clone before and after editing.

### 2.5 *In vitro* tri-lineage differentiation

iPSCs were aggregated into embryoid bodies using the hanging drop method and cultured for 14 days under spontaneous differentiation conditions. Differentiated outgrowths were replated onto gelatin-coated coverslips and analysed by immunofluorescence for β-III-tubulin (TUBB3; ectoderm), α-smooth muscle actin (α-SMA; mesoderm), and α-fetoprotein (AFP; endoderm), with DAPI counterstaining. Single-channel and merged images were acquired at 40× magnification under identical settings, and the proportion of marker-positive cells was quantified in three fields per lineage from each of three independent differentiations (nine fields per lineage), with isotype-only and undifferentiated-iPSC controls, using ImageJ **(Supplementary Table S5**).

### 2.6 Prime editing strategy and optimisation

The codon 8/9 (+G) insertion was corrected using PEmax, comprising Cas9-H840A nickase fused to an engineered M-MLV reverse transcriptase with optimised nuclear localisation signals (Anzalone et al., 2019; Chen et al., 2021). pegRNAs were designed with PEGIT (http://pegit.org, which ranks candidate spacers and enumerates PBS and RTT length combinations. Five candidate spacers were evaluated (scores 0.46–0.67), and the only candidate whose PAM is destroyed by the intended edit (GCATCTGACTCCTGAGGAGA; score 0.56) was selected so that the corrected allele cannot be re-engaged by the editor **(Supplementary Table S2**). The pegRNA was cloned into pU6-pegRNA-GG-acceptor (Addgene #132777) by BsaI Golden Gate assembly. PEmax was expressed from pCMV-PEmax (Addgene #174820) or, where MLH1dn was tested, from pCMV-PEmax-P2A-hMLH1dn (Addgene #174828), which co-expresses dominant-negative MLH1. Secondary nicking sgRNAs were synthesised as chemically modified oligonucleotides (2′-O-methyl at the three terminal nucleotides of each end plus 3′ phosphorothioate linkages; Integrated DNA Technologies). Plasmid identifiers and RRIDs are listed in Supplementary Table S4. Three parameters were varied independently. First, seven RTT lengths (16, 17, 18, 19, 20, 22, 24 nt) were tested with a fixed 13-nt PBS using PEmax alone. Second, four secondary nick positions were tested in a PE3 configuration using the optimal RTT length. Third, MLH1dn was co-expressed with and without the secondary nicking sgRNA. Each condition was performed in three independent electroporations (biological replicates), each harvested 72 h after electroporation.

#### Electroporation

iPSCs at 60–70% confluence was dissociated with TrypLE™ Express, resuspended in Buffer R, and electroporated on the Neon™ system with a 10 µL tip (1,100 V, 20 ms, 2 pulses) with ∼0.9 µg PEmax plasmid, 100 pmol pegRNA, and, where indicated, 50 pmol nicking sgRNA. Cells were plated on Matrigel in StemFlex™ with 10 µM Y-27632 and fed daily for 5–7 days.

### 2.7 Quantification of editing outcomes and off-target analysis

Single-cell selection was performed using the STRIPPER Micropipetter and confirmed through Sanger sequencing. For off-target analysis, deep amplicon sequencing was conducted. Paired-end reads were quality-filtered with fastp (Q≥20, minimum length 50 bp) and aligned to the human β-globin cluster reference NG_000007.3 using BWA-MEM; the entire cluster, not just *HBB*, was targeted to detect cross-amplification of HBD and HBG. Reads matched 99.0–99.9% to a 205 bp *HBB* exon 1 amplicon (NG_000007.3:70,551–70,755), covering codons 1–15 with the initiator ATG at position 70,595. Results were assigned to fragments at specific frequencies: bulk amplicon sequencing gave an intended-edit frequency of 8.4% and an indel frequency of 2.4%. All libraries were sequenced at a consistent depth (268,000 ± 3,000 read pairs, alignment rate 94.9%). Outcomes were quantified by direct haplotype counting for codons 1–15 and by CRISPResso2 (Clement et al., 2019), using the disease allele as reference and the corrected allele as the expected edited amplicon (20 bp window centered at −10, excluding 5 bp at each end).

### 2.8 Haematopoietic and erythroid differentiation

Three iPSC groups were differentiated in parallel: patient (non-edited) iPSCs, PEmax-treated (corrected, Homozygous) iPSCs, and control iPSCs. Embryoid bodies were formed on ultra-low-attachment plates. Mesoderm was induced from days 5–7 with BMP4 (50 ng/mL) and bFGF (50 ng/mL) in Stemline II™ medium. Haematopoietic specification was carried out between days 7–12 using VEGF, SCF, bFGF, TPO, IL-3, and insulin. From day 13 an extended formulation including EPO, holotransferrin and FLT3-L supported erythroid expansion and maturation. Complete cytokine concentrations and suppliers are listed in **Supplementary Table S3**. Cells were harvested on days 7, 14 and 22. Independent differentiations were initiated from separate iPSC passages on separate.

#### Flow cytometry

Cells were stained with CD71-FITC, CD235a-PE, CD34-APC and CD45-PerCP, each with matched isotype control. CD34/CD45 was assessed at day 12 and CD71/CD235a at days 7, 14 and 22. Samples were acquired on a BD FACSLyric cytometer (≥10,000 events per sample) and analysed in FlowJo v10 after exclusion of debris, doublets and dead cells; compensation was applied using single-stained controls. Erythroid data were analysed in two ways: **(i)** quadrant gating set on isotype controls, giving G−− (CD71⁻CD235a⁻), G+− (CD71⁺CD235a⁻), G++ (CD71⁺CD235a⁺) and G−+ (CD71⁻CD235a⁺), as in Figure 5a; and **(ii)** within the CD235a⁺ population, stratification into CD71-high, -intermediate and -low subsets to resolve stages of terminal maturation. Subset boundaries were set on the day-7 CD235a⁺ population and applied unchanged across all time points, conditions and replicates.

#### Globin gene expression and Protein expression

Quantitative RT-PCR was performed for *HBB*, *HBA1/HBA2*, *HBG1/HBG2*, *HBE1* and *HBD* with *GAPDH* normalisation (**Supplementary Table S1).** Data are presented as 2^−ΔΔCt^ relative to healthy-control erythroid cells for between-group comparison of each gene, and separately as 2^−ΔCt^ relative to *GAPDH* where absolute transcript abundance is compared. ΔCt values are not compared between genes because primer efficiencies differ between amplicons. Lysates were resolved by SDS-PAGE, transferred to PVDF, and probed with anti-β-globin (*HBB*) and anti-γ-globin (HBF) antibodies, with GAPDH as a loading control; signals were developed by enhanced chemiluminescence and imaged on a ChemiDoc system (Bio-Rad).

#### Haemoglobin HPLC

Cell pellets were lysed in hypotonic lysis buffer / Bio-Rad haemolysate reagent, clarified by centrifugation at 1,200 × g for 5 min, and analysed on the Bio-Rad VARIANT™ II Haemoglobin Testing System using the β-Thalassaemia Short Program, which resolves haemoglobin fractions by cation-exchange chromatography with detection at 415 nm (Bain et al., 2023). Peak identification, retention times, peak areas and area percentages were generated by the VARIANT™ II software using the manufacturer’s predefined retention-time windows. Peripheral blood from a healthy adult donor and from the patient was run in the same batch as assay control.

### 2.9 Statistical analysis

Data are presented as mean ± SD, with the number of independent biological replicates. Two-group comparisons used unpaired two-tailed Student’s t-tests. Comparisons across the three groups (patient, PEmax-treated, control) used one-way ANOVA at a single time point and two-way ANOVA for group and time point, each with Tukey’s post hoc test; time was treated as a repeated measure, with the same differentiation sampled serially. Where differentiations were matched by batch, comparisons were additionally analysed with differentiation batch as a blocking factor. In the prime editing optimization series, intended-edit versus indel frequency within a condition was compared by paired two-tailed t-test, and RTT lengths or nick positions by one-way ANOVA with Tukey’s post hoc test. P < 0.05 was considered statistically significant.

## 3. Results

### 3.1 Fibroblast isolation, reprogramming and pluripotency characterisation

Dermal fibroblasts obtained from the skin biopsy displayed the expected spindle-shaped mesenchymal morphology (**Figure 1a**). Sanger sequencing confirmed the homozygous frameshift variant c.27dupG (p.Ser10ValfsTer14) at codons 8/9 of *HBB* (**Figure 1b**). The cells stained positively for the mesenchymal markers type I collagen (Col1a1; **Figure 1d**), PDGFRα (**Figure 1e**) and vimentin (**Figure 1f**), and hematoxylin and eosin staining confirmed the expected cellular morphology (**Figure 1c**). Patient fibroblasts were reprogrammed with non-integrating Sendai virus vectors. Morphological changes were apparent 24 h after transduction, compact colonies emerged within one week, and colonies with characteristic pluripotent morphology and sharply demarcated borders were established by three weeks (**Figure 2A**). G-banding confirmed a normal 46, XY karyotype (**Figure 2B**). Immunofluorescence demonstrated nuclear expression of the core pluripotency transcription factors OCT4, SOX2 and NANOG, and surface expression of TRA-1-60 (**Figure 2C**). Single-channel images are presented alongside merged overlays so that nuclear versus surface localization can be assessed independently for each marker.

**Figure 1.**
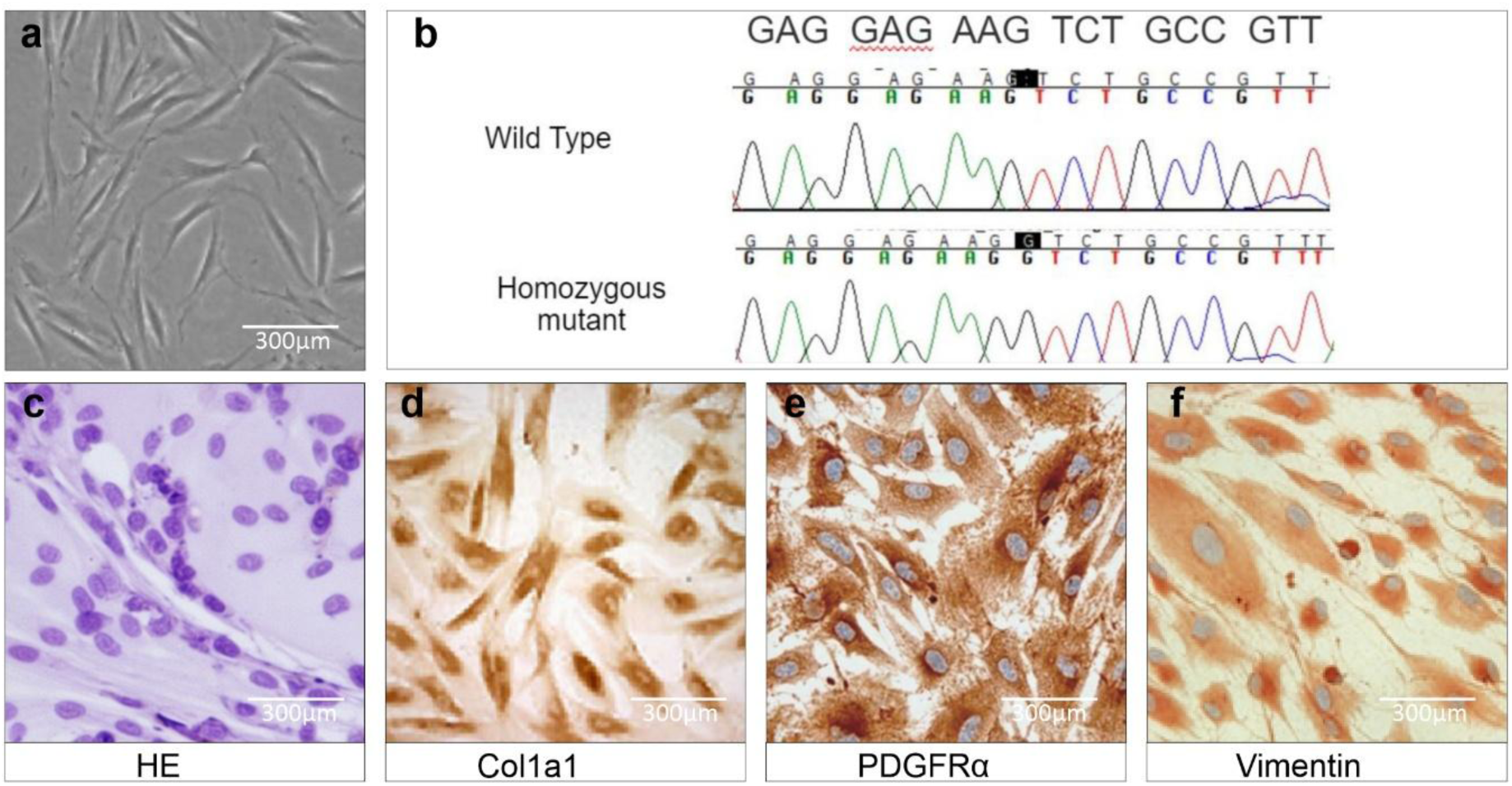
Isolation and characterisation of patient-derived dermal fibroblasts. **(a)** Phase-contrast micrograph of primary fibroblasts isolated from a β-thalassaemia patient, showing typical spindle-shaped morphology (scale bar, 300 µm). **(b)** Sanger sequencing chromatograms confirming the homozygous HBB codon 8/9 (+G) mutation (c.27dupG) in patient-derived fibroblasts relative to a wild-type control. **(c–f)** Fibroblast identity confirmed by **(c)** H&E staining and immunocytochemistry for **(d)** Col1a1, **(e)** PDGFRα, and **(f)** Vimentin (scale bars, 300 µm).

**Figure 2.**
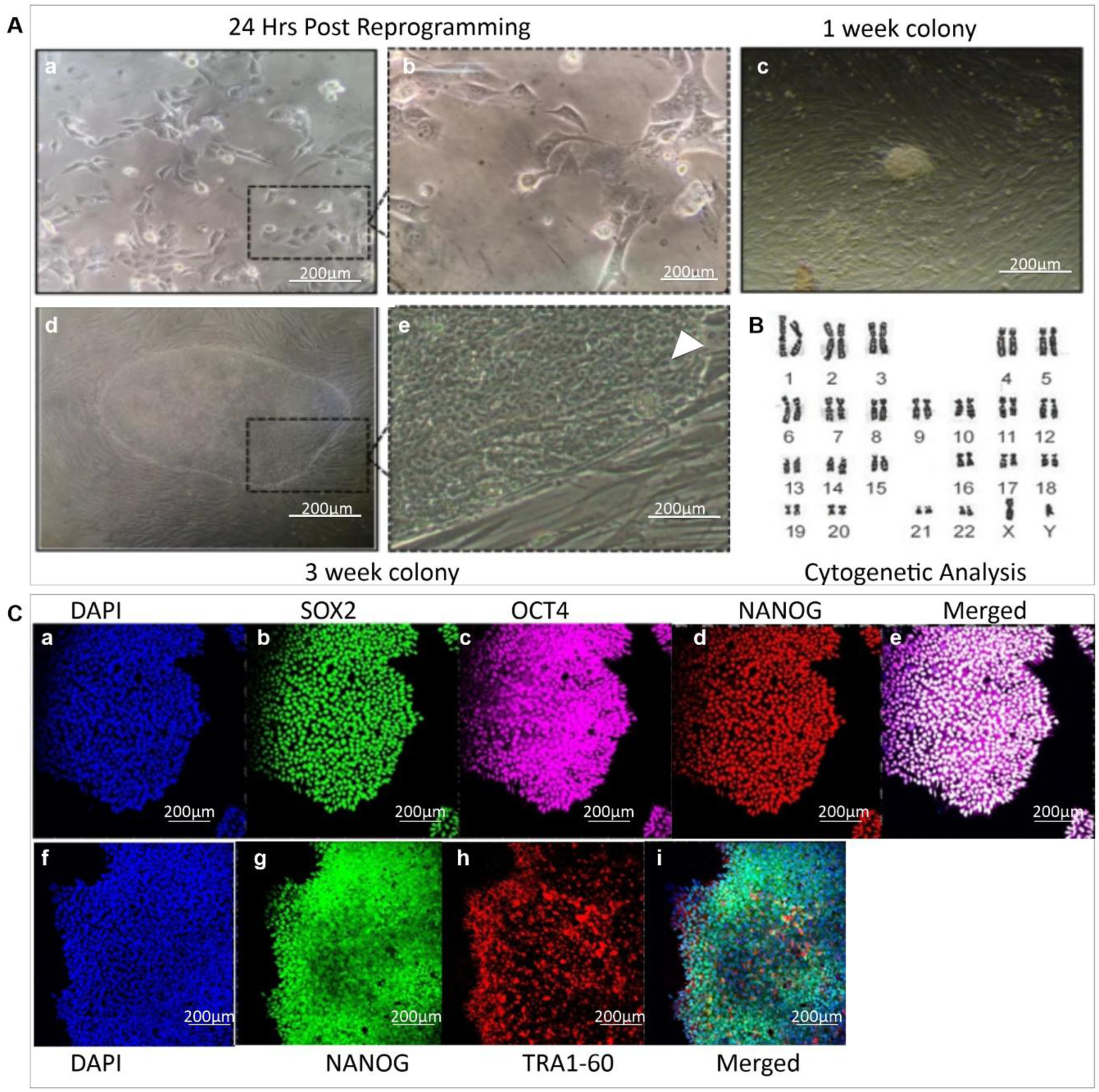
Reprogramming of thalassaemic fibroblasts into iPSCs. **(A)** Phase-contrast time-course of reprogramming: partially reprogrammed cells at 24 h post-transduction (a, low magnification; b, high magnification), an emerging colony at 1 week (c), and a mature iPSC-like colony at 3 weeks (d, e) (scale bars, 200 µm). **(B)** G-banded karyotype (46,XY) showing a normal diploid chromosome complement with no gross structural or numerical abnormalities. **(C)** Immunofluorescence staining for core pluripotency markers: (a–e) DAPI, SOX2, OCT4, NANOG, and merged; (f–i) DAPI, NANOG, TRA-1-60, and merged (scale bars, 200 µm)

### 3.2 Functional pluripotency, endogenous gene expression and Sendai vector clearance

Embryoid bodies derived from patient iPSCs generated derivatives of all three germ layers, identified by TUBB3 (ectoderm), α-SMA (mesoderm) and AFP (endoderm) immunofluorescence, each with the expected cytoplasmic or filamentous distribution (Figure 3a). Marker-positive cells represented 22.7%, 26.2% and 25.9% of DAPI⁺ nuclei for TUBB3, AFP and α-SMA, respectively (mean of three fields in each of three independent differentiations; **Supplementary Table S5**), compared with ≤1.8% in isotype-only and undifferentiated-iPSC controls. Quantitative RT-PCR of six endogenous pluripotency genes showed that patient iPSCs expressed *OCT4*, *SOX2*, *NANOG*, *LIN28*, *REX1* and *DNMT3B* at approximately 72–94% of the levels in human ESCs, while parental fibroblasts showed negligible expression (**Figure 3b)**. As a percentage of ESC levels, iPSC values were approximately 94%, 87% and 91% for *OCT4*, *SOX2* and *NANOG*, and approximately 76%, 83% and 72% for *LIN28*, *REX1* and *DNMT3B*, with overlapping error bars for several genes; no statistical comparison was performed. The lines satisfied the functional criteria assessed, including tri-lineage differentiation and a normal karyotype. Sendai vector clearance was assessed by TaqMan RT-PCR for the SeV backbone and the KOS, *KLF4* and *c-MYC* transgenes **(Figure 3c**). Transduced fibroblasts showed strong amplification of all four targets. At passage 2, all four remained detectable, whereas at passage 15 no amplification was observed within 40 cycles for any of the four assays, while the endogenous control amplified normally, meeting the pre-specified criterion for a vector-free line.

**Figure 3.**
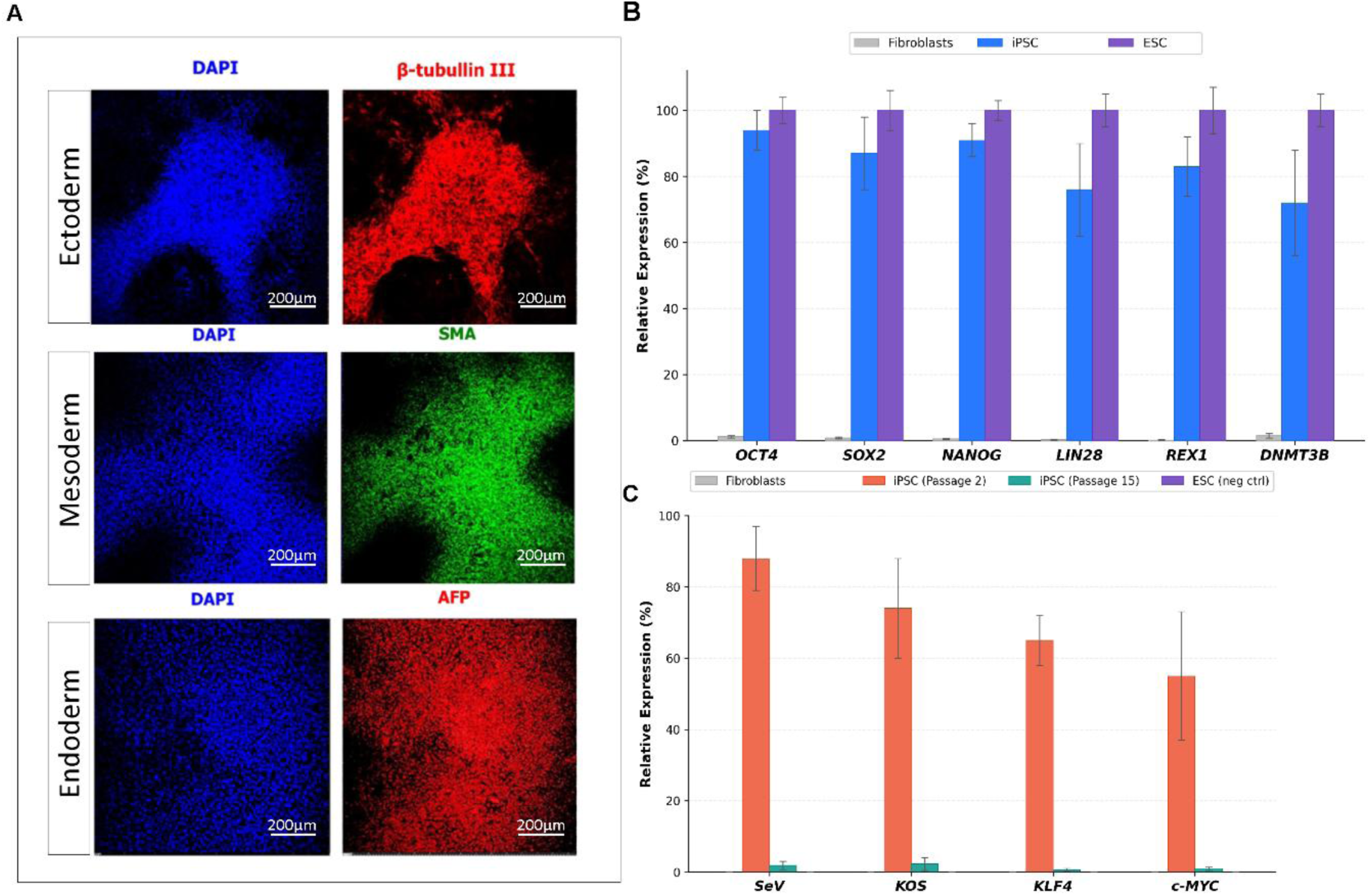
Pluripotency validation and vector silencing. **(a)** Trilineage differentiation potential confirmed by immunostaining for ectoderm (β-tubulin III), mesoderm (SMA), and endoderm (AFP) markers, with DAPI counterstain (scale bars, 200 µm). **(b)** Relative expression (%) of endogenous pluripotency genes (*OCT4, SOX2, NANOG, LIN28, REX1, DNMT3B*) in parental fibroblasts, iPSCs, and an ESC reference line. **(c)** Relative expression (%) of Sendai virus vector/transgene sequences (*SeV, KOS, KLF4, c-MYC*) in fibroblasts and ESCs (negative controls) versus iPSCs at passage 2 and passage 15, confirming vector clearance with continued passaging.

### 3.3 Systematic optimisation of prime editing at the *HBB* codon 8/9 locus

**(a)** Five candidate pegRNA spacers were evaluated computationally (**Supplementary Table S2**) and the spacer GCATCTGACTCCTGAGGAGA was selected because it was the only candidate whose PAM is destroyed by the intended edit, thereby preventing re-engagement of the corrected allele (**Figure 4a**). Editing outcomes were quantified by targeted amplicon deep sequencing at a median depth of 268,000 reads per sample across three independent electroporations per condition. Varying the RTT length from 16 to 24 nt against a fixed 13-nt PBS produced length-dependent differences in the intended-edit frequency, which was highest at 17nt and declined at both shorter and longer extensions (**Figure 4c**). Introduction of a secondary nick in the PE3 configuration at four positions within ±70 bp of the pegRNA-directed nick showed clear position dependence, with the +68 bp position yielding the highest intended-edit frequency (8.4%); as expected for PE3, the gain in intended editing was accompanied by an increase in indel frequency relative to PE2 **(Figure 4d**). Co-expression of MLH1dn had only a minor effect. Under the combined optimal condition (RTT 17 nt, PBS 13 nt, nick at +68 bp, MLH1dn co-expression), bulk amplicon sequencing gave an intended-edit frequency of 8.4% and an indel frequency of 2.4%. (**Supplementary Figure S1**). 96 clones were analyzed for the editing. This yielded 3 homozygous corrected clones (3.1%), 11 heterozygous (11.5%), 77 unedited (80.2%), and 5 with indels (5.2%), indicating an overall editing efficiency of 19.8% (**c**) show normal karyotype after editing. (**Figure 4e; Supplementary Figure S1**).

**Figure 4.**
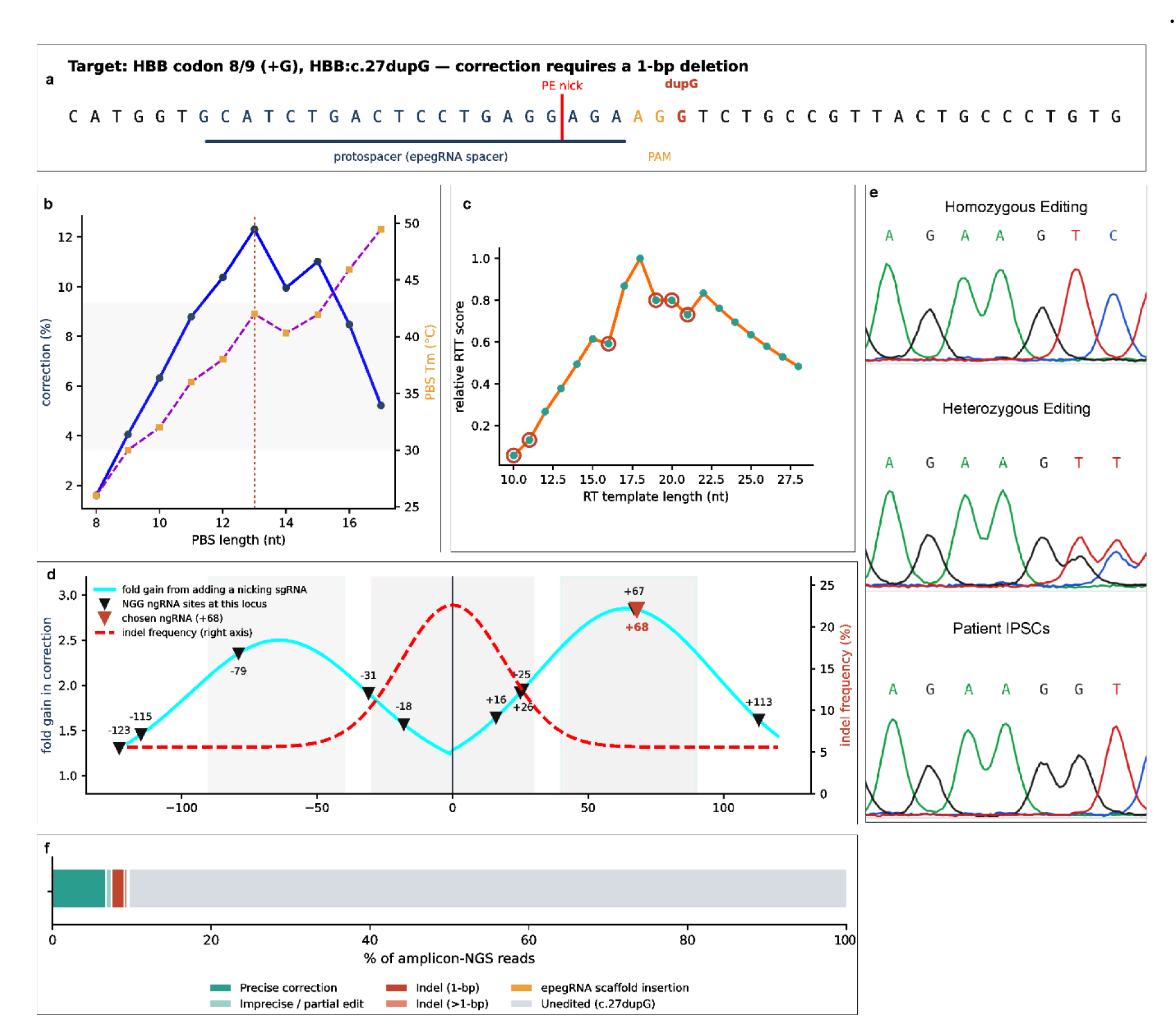
Prime editing strategy for correction of the HBB c.27dupG mutation(a) Schematic of the target locus showing the pegRNA protospacer, PAM, PE nick site, and the duplicated G (dupG) requiring a 1-bp deletion for correction. **(b)** PBS length optimization: correction efficiency (%, left axis) and predicted PBS melting temperature (°C, right axis) across PBS lengths. **(c)** RTT length optimization based on relative RT template (RTT) score. **(d)** Nicking sgRNA screen: fold-gain in correction efficiency and indel frequency (%) plotted by distance from the pegRNA nick site, with the selected +68 nicking sgRNA indicated. **(e)** Sanger chromatograms confirming homozygous correction, heterozygous correction, and the uncorrected patient sequence. **(f)** NGS-based quantification of editing outcomes at the target locus.

### 3.4 Validation of the amplicon-sequencing quantification pipeline on libraries

To validate the quantification pipeline, we analysed the amplicon libraries. All five outcome classes were resolved as distinct haplotypes: the corrected allele carried GCA at codon 10 and translated to wild-type MVHLTPEEKSAVTAL, while the partially corrected allele retained the frameshift. Direct haplotype counting returned the unedited disease allele at 84.9% ± 0.09%, perfect correction at 8.6 ± 0.02%, partial correction at 2.4 ± 0.01%, indels at 1.10 ± 0.01% and scaffold insertion at 0.37% of spanning reads ; recovery was uniform at 91.4–92.5% across all five classes, the shortfall reflecting inclusion of the 4.2–4.6% of fragments whose sequencing errors disrupted the target context in the denominator rather than class-specific detection bias. Across ten PE-treated replicates at uniform depth, CRISPResso2 independently recovered the precise correction frequency at 8.4 ± 0.03%, while the mock-treated and wild-type control arms returned 0.30% and 0.33% respectively, cleanly separating background misassignment from genuine correction. Raw amplicon indel counts (∼4.4%) were dominated by background also seen in mock controls (∼4.3%); the CRISPResso2 windowed indel frequency was 2.4%

### 3.5 Haematopoietic and erythroid differentiation of patient, corrected and control iPSCs

Erythroid differentiation of patient-derived, PEmax-treated and control iPSCs was assessed by CD71 and CD235a expression at days 7, 14 and 22 (**Figure 5a**) and by colony-forming assays over the same time course (**Figure 5b**). At day 7, cultures from all three conditions were predominantly CD71⁻CD235a⁻ (patient 87.2%, PEmax 80.6%, control 79.9%), and CD235a⁺ cells (CD71⁻ plus CD71⁺) made up 8.4%, 10.9% and 11.6% of cells, respectively. By day 14 the patient culture had progressed further along the erythroid axis. CD71⁺CD235a⁺ cells accounted for 65.8% of patient cells, compared with 39.1% in PEmax-treated and 49.8% in control cultures, while the double-negative fraction fell to 9.1% versus 37.9% and 27.2%. At day 22 the total CD235a⁺ fraction was similar across conditions (patient 93.9%, PEmax 91.9%, control 93.6%), but its composition differed. The CD71⁺CD235a⁺ fraction was lower in patient cells (82.8%) than in PEmax-treated (91.1%) or control (93.1%) cells, and the CD71⁻CD235a⁺ fraction was higher (11.0% versus 0.8% and 0.5%). At day 22, the patient culture also showed a diffuse CD71 distribution, whereas PEmax-treated and control cells formed a compact CD71⁺CD235a⁺ population. Colony-forming assays showed a parallel pattern (Figure 5b). CFU-E numbers were lower in patient cultures than in control cultures at day 7 (P < 0.05) and than in both PEmax-treated and control cultures at day 14 (about 195 versus 365 and 375 colonies per 10⁴ cells; P < 0.01).

**Figure 5.**
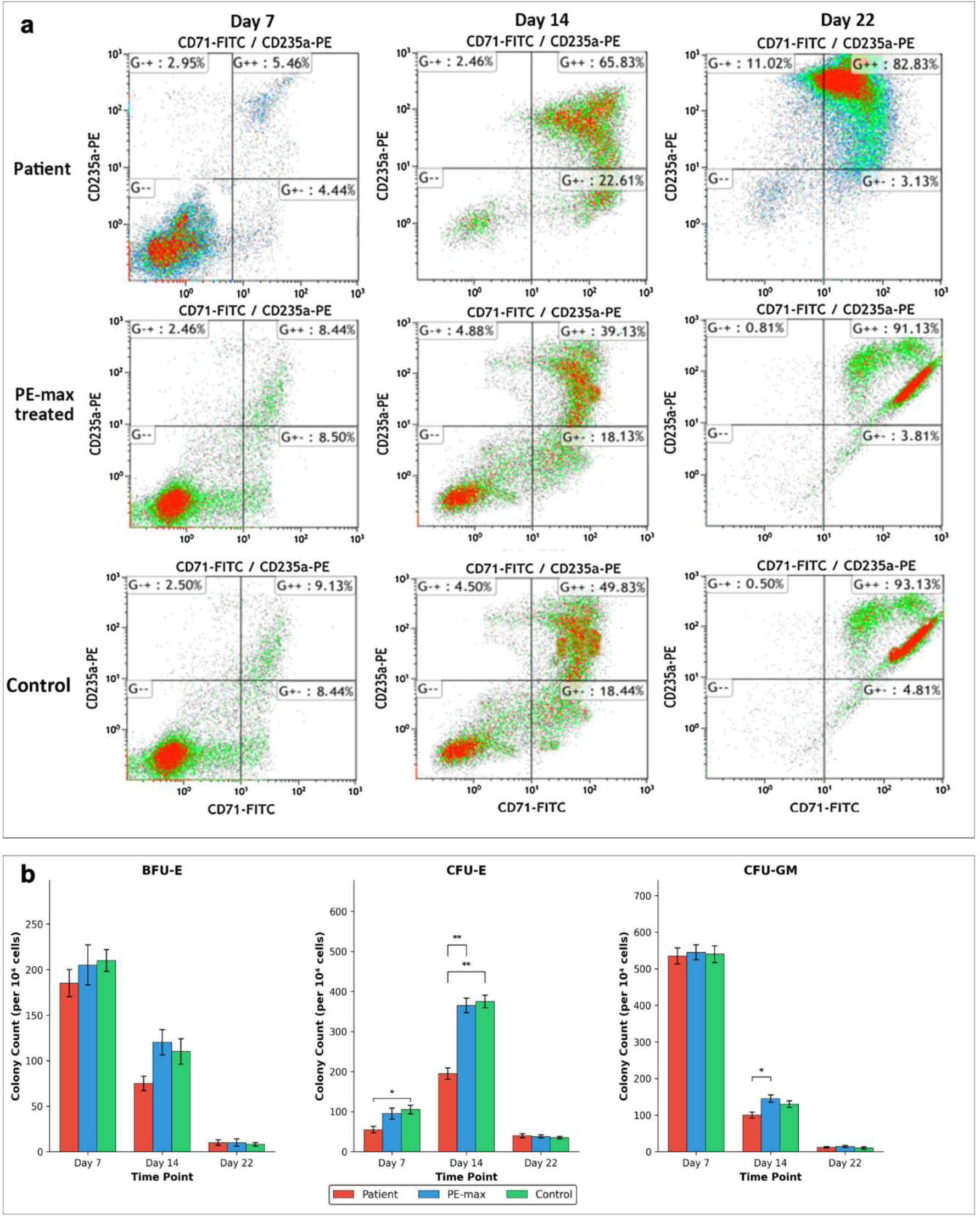
Erythroid differentiation of corrected iPSCs. **(a)** Flow cytometry analysis of CD71/CD235a expression at days 7, 14, and 22 of erythroid differentiation in patient, PEmax-treated, and control iPSC lines, showing progressive maturation (single representative plot per condition; quadrant labels as defined in Methods 2.8). **(b)** Colony-forming assays quantifying BFU-E, CFU-E, and CFU-GM numbers (per 10⁴ cells) across the same time points and groups (*P < 0.05, **P < 0.01).

BFU-E counts did not differ significantly between conditions, and CFU-GM was lower in patient cultures only on day 14 (∼100 versus ∼145 in PEmax). Colony numbers of all types converged at day 22. Across both readouts, PEmax-treated cells tracked the control more closely than the patient cells.

### 3.6 Restoration of β-globin transcript and protein, and detection of an HbA fraction

To test whether prime editing corrected the *HBB* codon 8/9 frameshift, we first quantified globin transcripts by RT-qPCR **(Figure 6a)**. *HBB* mRNA was almost undetectable in patient iPSC-derived cells (∼0.05-fold relative to control iPSCs, P < 0.01) and in patient peripheral blood (PB) cells. PEmax treatment restored HBB expression to ∼4-fold that of control iPSC-derived cells (n = 3; patient vs. treated, P < 0.01; control vs. treated, P < 0.05); expression above the control level may reflect differences in erythroid maturation between lines rather than supraphysiological expression and is not interpreted further. Levels remained far below those of adult PB control cells (∼1,000-fold), as expected for cells with a fetal-type globin programme. *HBG1/HBG2* (γ-globin) and *HBE1* (ε-globin) transcripts were abundant in all iPSC-derived groups and near-absent in PB cells. Neither changed significantly with editing, although *HBG1/HBG2* showed a non-significant upward trend. At the protein level (Figure **6b**), HBB was not detectable in patient iPSC-derived cells or PB patient cells. Edited cells showed a faint HBB band comparable to that of control iPSCs, whereas PB control cells showed a strong band. These three readouts are not expected to scale with one another: RT-qPCR reports relative transcript fold-change against a near-zero patient baseline (so a small absolute increase produces a large fold-value), western blotting reports steady-state β-globin protein semi-quantitatively, and HPLC reports the assembled α/β tetramer as a percentage of total haemoglobin. A faint β-globin band and a ∼6% tetramer fraction is therefore consistent with a low absolute level of correctly assembled adult hemoglobin against a predominantly fetal/embryonic background, rather than being contradictory. HBF was the predominant globin in all iPSC-derived samples. Haemoglobin HPLC (**Figure 6c**) showed the same pattern .A cation-exchange fraction eluting in the HbA retention-time window was undetectable in-patient iPSC-derived cells and rose to ∼6% after editing (P < 0.001), not significantly different from control iPSCs (∼6%). This fraction is consistent with, but not confirmed as, HbA: it eluted at a retention time differing from adult HbA in the peripheral-blood control and close to the HbA2 window, and orthogonal confirmation by intact-globin mass spectrometry or co-elution with an adult HbA standard was not performed.

**Figure 6.**
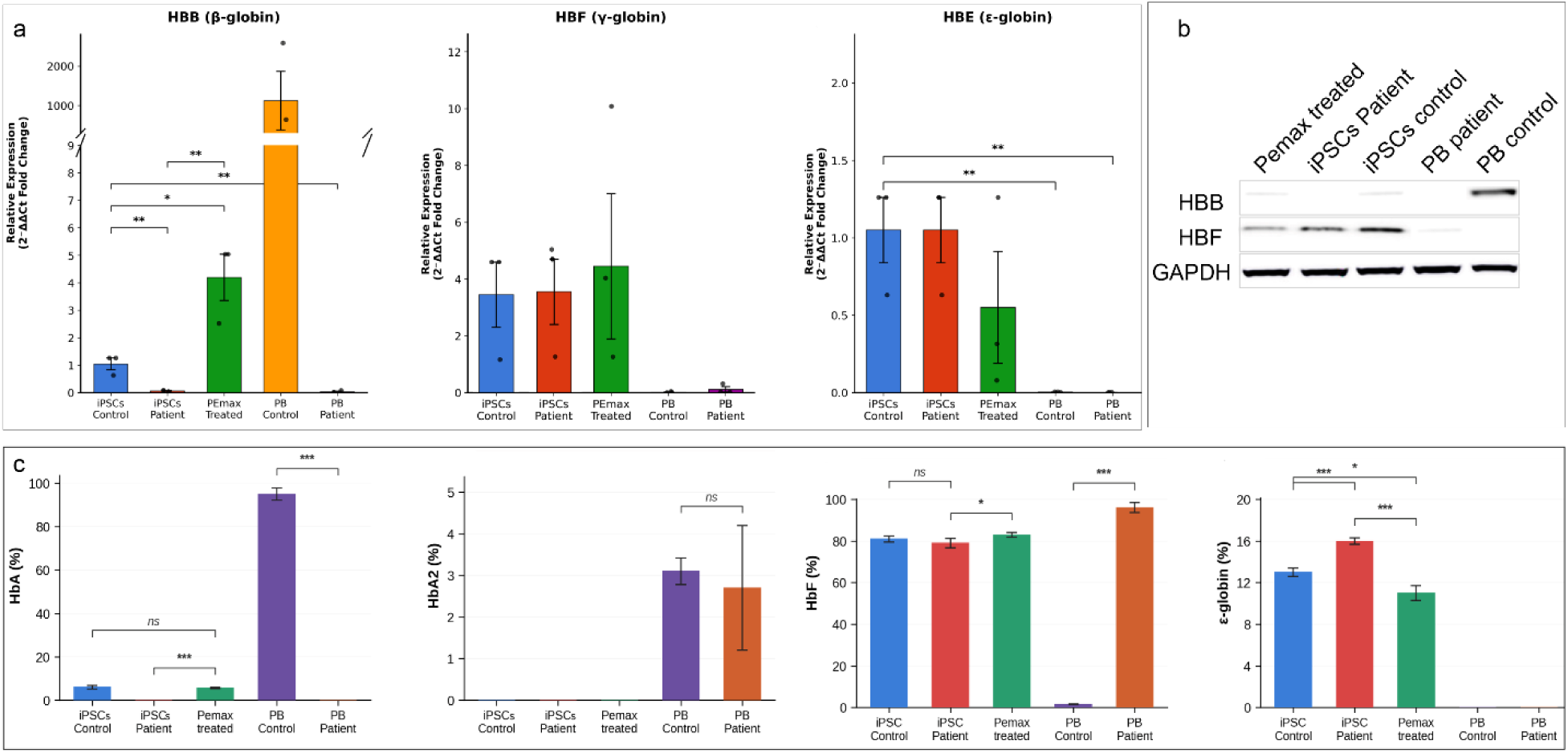
Restoration of globin expression following gene correction. (**a**) qRT-PCR analysis of *HBB* (β-globin), *HBG1/HBG2* (γ-globin) and *HBE1* (ε-globin) relative expression in iPSC-derived erythroid cells and peripheral blood (PB) controls/patients (*P < 0.05, **P < 0.01). **(b)** Western blot of HBB and HBF protein (GAPDH loading control) across PEmax-treated, patient, and control iPSC-derived erythroid cells and PB samples. **(c)** HPLC-quantified hemoglobin fractions (HbA, HbA2, HbF, ε-globin, %) showing an HbA fraction in PEmax-treated cells that is undetectable in-patient cells, and a lower ε-globin fraction after correction (**P < 0.001, *P < 0.05; ns, not significant.

By comparison, PB control cells contained ∼95% HbA. HbA2 was detected only in PB samples. HBF remained dominant in all iPSC-derived cells (∼79–83%), with a small increase after editing (P < 0.05). The fraction assigned to embryonic ε-globin-containing haemoglobin was highest in patient cells (∼16%) and decreased after editing (∼11%), slightly below that of control iPSC-derived cells (∼13%; P < 0.05). Together, these data indicate that editing restored HBB transcript and detectable β-globin protein and yielded a candidate HbA fraction comparable to that of control iPSC-derived cells (∼6%), while absolute adult haemoglobin remained low in the fetal/embryonic globin programme of iPSC-derived erythroid cells.

## 4. Discussion

We combined derivation of integration-free patient iPSCs, locus-specific optimization of PEmax prime editing and multi-level erythroid readouts to test correction of the HBB codon 8/9 (+G) frameshift. Editing was followed by restoration of *HBB* transcript and detectable β-globin protein, and by an HPLC fraction assigned as HbA that was undetectable in uncorrected cells, within an erythroid programme that remained predominantly fetal and embryonic.

The patient-derived lines met standard criteria for pluripotency: a normal 46,XY karyotype (Figure 2B), nuclear OCT4, SOX2 and NANOG (Figure 2C), derivatives of all three germ layers (Figure 3a) and no detectable Sendai transcripts by passage 15 (Figure 3c). Dermal fibroblasts proved a practical starting population (Mali et al., 2008), integration-free Sendai reprogramming avoids transgene integration (Beers et al., 2015), and quantitative TaqMan assays give a standardised measure of vector clearance. Endogenous pluripotency genes reached approximately 72–94% of ESC levels, with *LIN28*, *REX1* and *DNMT3B* at the lower end; because error bars overlapped and no statistical test was applied, we do not attribute this to incomplete reprogramming or epigenetic memory. Embryoid-body outgrowths contained 23–26% lineage-marker-positive nuclei **(Supplementary Table S5**), which shows lineage potential rather than differentiation efficiency.

Editing efficiency depended on pegRNA design. Only one of the five candidate spacers destroys its own PAM on correction, because deleting the extra G converts the PAM from AGG to AGT; this prevents re-nicking of the corrected allele, a design advantage that becomes more important as efficiency rises. We selected it although two alternatives had higher predicted scores (**Supplementary Table S2).** A secondary nick at +68 increased intended edits at the cost of more indels, the expected PE3 trade-off (Anzalone et al., 2019), whereas MLH1dn co-expression added little, unlike the substantial gains reported in other settings (Chen et al., 2021). Delivery, cell type, edit type and modest overall efficiency may all contribute, but we did not test this. An intended-edit frequency of about 8.4 % is modest; structured 3′ motifs that protect pegRNAs (epegRNAs; Nelson et al., 2022) could be tested at this locus. The bulk intended-edit frequency (8.4% of alleles at 72 h) and the clonal correction rate (19.8% of clones carrying ≥1 corrected allele) are not directly comparable, as a clone scores as corrected if either allele is edited; a clone-level frequency exceeding the allele-level frequency is therefore expected, consistent with the allele-to-clone relationships reported for prime editing in human pluripotent stem cells (Sürün et al., 2022; Kanno et al., 2025). The recovery of biallelically corrected clones (3.1%) is at the higher end of what is typically observed when editing from a homozygous starting genotype, where homozygous conversion is generally the rarer outcome (Kanno et al., 2025; Li et al., 2024); we attribute this to the limited clone number (n = 96) and possible enrichment of edited cells during single-cell outgrowth, rather than to supraphysiological editing, and note it as a limitation pending confirmation in additional clones

In erythroid culture, PEmax-treated cells resembled control cells more closely than patient cells in CD71/CD235a kinetics and in CFU-E and CFU-GM output. Patient cells were unexpectedly ahead at day 14 (65.8% CD71⁺CD235a⁺ versus 39.1% and 49.8%) but retained a CD71⁻CD235a⁺ subset at day 22 (11.0% versus 0.8% and 0.5%); these flow data are single representative plots. In β-thalassaemia, unpaired α-globin chains precipitate in erythroid precursors and drive oxidative damage, apoptosis, and ineffective erythropoiesis (Rivella, 2012), so a corrective effect on terminal erythroid maturation is mechanistically plausible. We did not, however, measure α/β chain ratios, α-globin precipitation, reactive oxygen species or apoptosis in these cultures; therefore, we describe an association rather than a demonstrated mechanism. A further consideration specific to this model is that the differentiated cells express predominantly γ- and ε-globin, so the α-chain excess that drives the disease phenotype in adult erythropoiesis is substantially buffered by fetal and embryonic β-like chains. The magnitude of any maturation defect attributable to β⁰ genotype is correspondingly expected to be smaller in iPSC-derived erythroid cells than in patient bone marrow, which should be borne in mind when interpreting differences between groups as evidence of phenotypic rescue.

The globin data are best read as correction of the genetic lesion with partial, not complete, restoration of an adult haemoglobin programme. *HBB* transcript in PEmax-treated cells exceeded that in control iPSC-derived cells (about 4-fold, n = 3), which points to differences in maturation stage or normalisation between lines rather than supraphysiological expression; consistent with this, the fold-change in transcript, the faint protein band and the ∼6% HPLC fraction reflect different quantities (relative mRNA, steady-state protein, assembled tetramer) and are not expected to be numerically concordant. This fetal/embryonic bias is a well-established property of erythroid cells generated from human pluripotent stem cells, which typically recapitulate primitive or fetal-definitive rather than adult-definitive erythropoiesis and rarely complete the γ-to-β switch under standard *in vitro* conditions (Dias et al., 2011; Lapillonne et al., 2010; Kobari et al., 2012). Forced expression of KLF1 and BCL11A-XL can induce adult-level β-globin in such cells (Trakarnsanga et al., 2014), and serum-free, feeder-free small-molecule protocols have improved erythroid output and maturation (Olivier et al., 2016). Corrected sickle-cell iPSCs have likewise yielded erythroid cells with adult β-globin protein (Huang et al., 2015).

Earlier iPSC-based correction studies for β-thalassemia used nuclease-induced homology-directed repair. Xie et al. (2014) corrected the −28 (A>G) and codon 41/42 mutations with Cas9 and piggyBac, Xu et al. (2015) targeted IVS2-654, and Niu et al. (2016), Yang et al. (2016) and Liu et al. (2017) corrected codon 41/42 by CRISPR/Cas9-mediated homology-directed repair; Song et al. (2015) reported improved haematopoietic differentiation of gene-corrected β-thalassaemia iPSCs. These approaches depend on double-strand breaks, with attendant risks of indel formation, large deletions, translocations and p53-mediated toxicity (Haapaniemi et al., 2018). Prime editing avoids double-strand-break intermediates and exogenous donor templates (Anzalone et al., 2019) and has been applied to *HBB* in erythroid progenitor cells (Zhang et al., 2022) and, ex vivo, to sickle-cell HSPCs with engraftment in mice (Everette et al., 2023). To our knowledge, its use for c.27dup (p.Ser10ValfsTer14) in patient iPSCs with an erythroid readout has not been reported.

Direct editing of haematopoietic stem and progenitor cells remains the more immediate clinical route (Dever et al., 2016; Hoban et al., 2016; Yu et al., 2016; Cottle et al., 2016). The iPSC platform is complementary, offering clonal isolation, including allele-specific editing (Smith et al., 2015), comprehensive genomic quality control and unlimited expansion for systematic optimisation of editing strategies. Exagamglogene autotemcel has shown that *BCL11A*-mediated HbF induction can achieve transfusion independence (Frangoul et al., 2021; Locatelli et al., 2024) without addressing the underlying *HBB* defect, and other approaches recreate hereditary-persistence-of-fetal-haemoglobin promoter variants (Traxler et al., 2016). Direct correction and HbF induction target different mechanisms and are not mutually exclusive; whether their combination would be additive remains untested.

Several limitations should be emphasized. The study rests on a single patient-derived iPSC line and a limited number of corrected clones, so findings require confirmation across additional donors carrying this variant. Flow cytometry is shown as single representative plots, RT-qPCR has three points per group, so the reported P values should be interpreted cautiously. The peaks assigned as HbA and embryonic hemoglobin in **Supplementary Figure S3** eluted at retention times that differ from adult HbA in the peripheral-blood control and near the HbA2 window, and HPLC of small-scale cultures is affected by low hemoglobin content and co-eluting peaks; orthogonal confirmation is needed. Off-target assessment was limited to 35 in-silico-predicted sites examined by Sanger sequencing and deep amplicon sequencing, which cannot detect structural variants; unbiased methods such as GUIDE-seq (Tsai et al., 2015) or CIRCLE-seq (Tsai et al., 2017), whole-genome sequencing were not applied. Editing was performed in iPSCs rather than hematopoietic stem and progenitor cells, iPSC-derived hematopoietic cells have not been shown to reconstitute long-term hematopoiesis, the differentiated cells retain a fetal/embryonic globin programme irrespective of genotype, no in vivo engraftment was performed.

## 5. Conclusions

In patient-derived iPSCs homozygous for *HBB* c.27dupG, PEmax prime editing with a PAM-disrupting pegRNA and a +68 nicking sgRNA achieved an intended-edit frequency of 8.4% in the bulk population and yielded corrected cells. Erythroid cells derived from them restored *HBB* transcript and β-globin protein, contained an HPLC fraction consistent with HbA that was absent from uncorrected cells, and matured more like control cells than patient cells. All iPSC-derived cells remained dominated by fetal and embryonic globins, so these data show genetic correction with partial restoration of adult hemoglobin synthesis rather than a functional cure. Confirmation across donors and clones, in hematopoietic stem and progenitor cells with engraftment, and with comprehensive off-target and structural-variant analysis is needed before clinical translation. The optimization workflow described here (spacer choice, extension length, nick position, MLH1dn) may be applicable to other *HBB* variants.

## Declarations Funding

This work was supported by Wellcome Leap (project 53452).

## Conflicts of interest

The authors declare no competing financial or non-financial interests.

## Ethics approval

Approved by the Aga Khan University Ethics Review Committee (2022-6532-22590). Written informed consent was obtained from all participants in accordance with the Declaration of Helsinki.

## Availability of data and materials

Data other than the main and supplementary files are available on request

## Author contributions

I.H. conceived and designed the study, performed computational pegRNA design, cloning, prime editing optimization, cell culture, hematopoietic differentiation and data analysis, and wrote the original draft. K.M. contributed to computational design, cloning, prime editing experiments, cell culture and differentiation. S.F. assisted with prime editing optimization, cloning, cell culture and differentiation. A.A. contributed to computational design, prime editing optimization, cell culture and differentiation. A.F. and H.Z. performed iPSC derivation, colony selection, pluripotency characterization and tri-lineage differentiation. M.J. and F.R. performed the skin biopsy, fibroblast isolation and culture, karyotyping and Sanger sequencing for genotyping and off-target validation. M.J. contributed to data curation. S.K. provided clinical guidance, supervised the study and revised the manuscript. F.S. supervised the study, contributed to protocol optimization and revised the manuscript. A.A.M. conceptualized and supervised the study, acquired funding, optimized protocols and edited the final manuscript. All authors read and approved of the final manuscript.

## Acknowledgements

The authors thank the Centre for Regenerative Medicine and Stem Cell Research at Aga Khan University for infrastructure and institutional support; Dr Syed Ather Enam, Director of the Centre, for institutional backing; Dr Azhar Hussain, Laboratory Director, for technical oversight; and Ms Sadia, Laboratory Manager, for day-to-day laboratory support.

## Supplementary material

**Supplementary Table S1.** Primer sequences for qRT-PCR, globin gene expression, *HBB* genotyping and amplicon sequencing.

**Supplementary Table S2.** pegRNAs, including candidate spacers with scores, nicking sgRNA candidates and the full PBS/RTT length matrix.

**Supplementary Table S3.** Cytokines, growth factors, media and suppliers for haematopoietic differentiation.

**Supplementary Table S4.** Plasmids used with Addgene identifiers and RRIDs.

**Supplementary Table S5.** Quantification of tri-lineage marker-positive cells, with isotype-only and undifferentiated-iPSC controls.

**Supplementary Table S6.** Targeted amplicon sequencing summary: read depth, intended edit, indel, edited and wild-type allele frequencies for every condition and replicate.

**Supplementary Table S7.** Predicted off-target sites and Sanger sequencing results.

---

**Figure S1:**
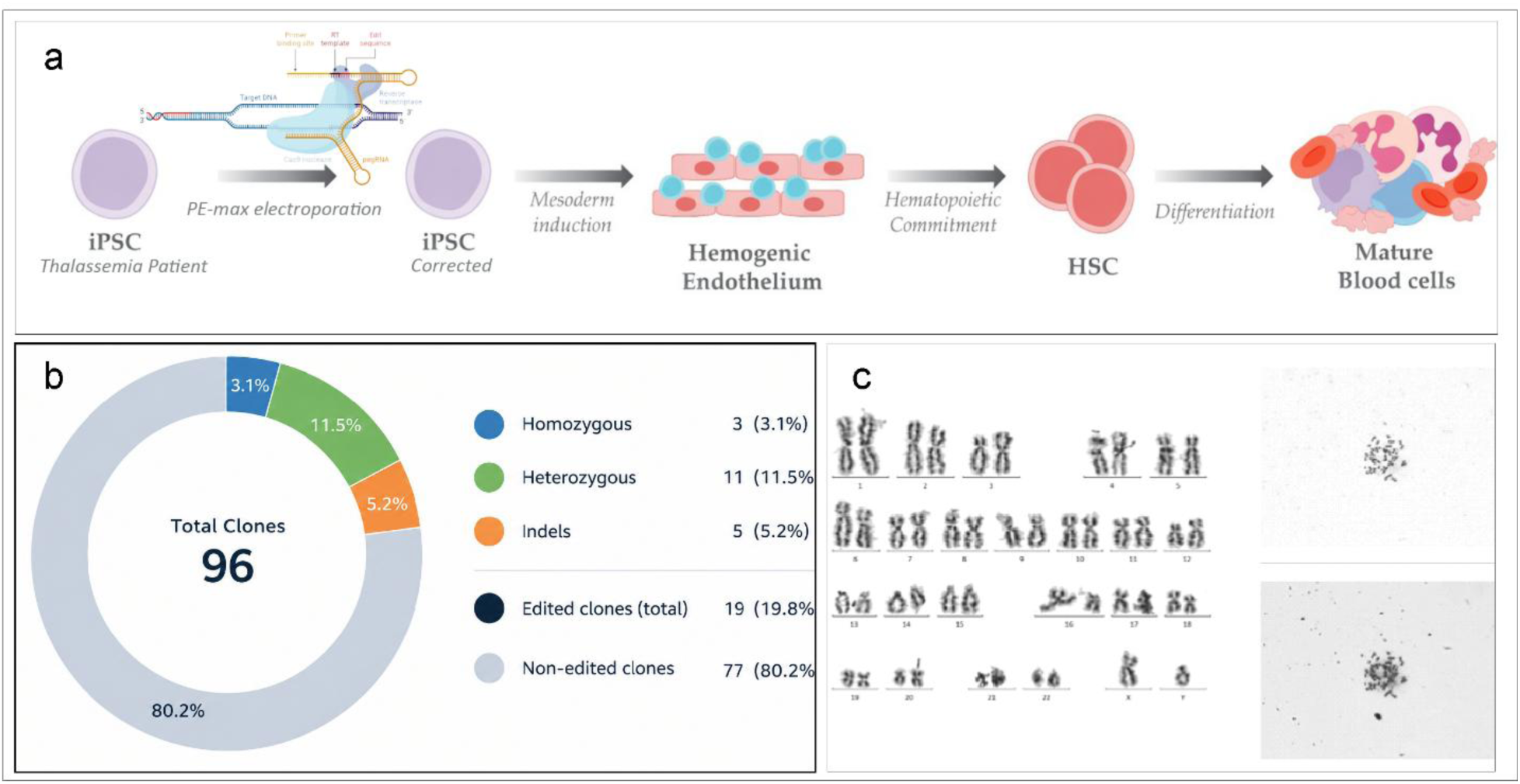
Clonal validation. **(b)** Depicts the overall methodology **(b)** show the validation of editing by sanger sequencing. 96 clones were analyzed by Sanger sequencing (Figure 5c–d). This yielded 3 homozygous corrected clones (3.1%), 11 heterozygous (11.5%), 77 unedited (80.2%), and 5 with indels (5.2%), indicating an overall editing efficiency of 19.8% (c) show normal karyotype after editing.

**Supplementary Figure S1.**
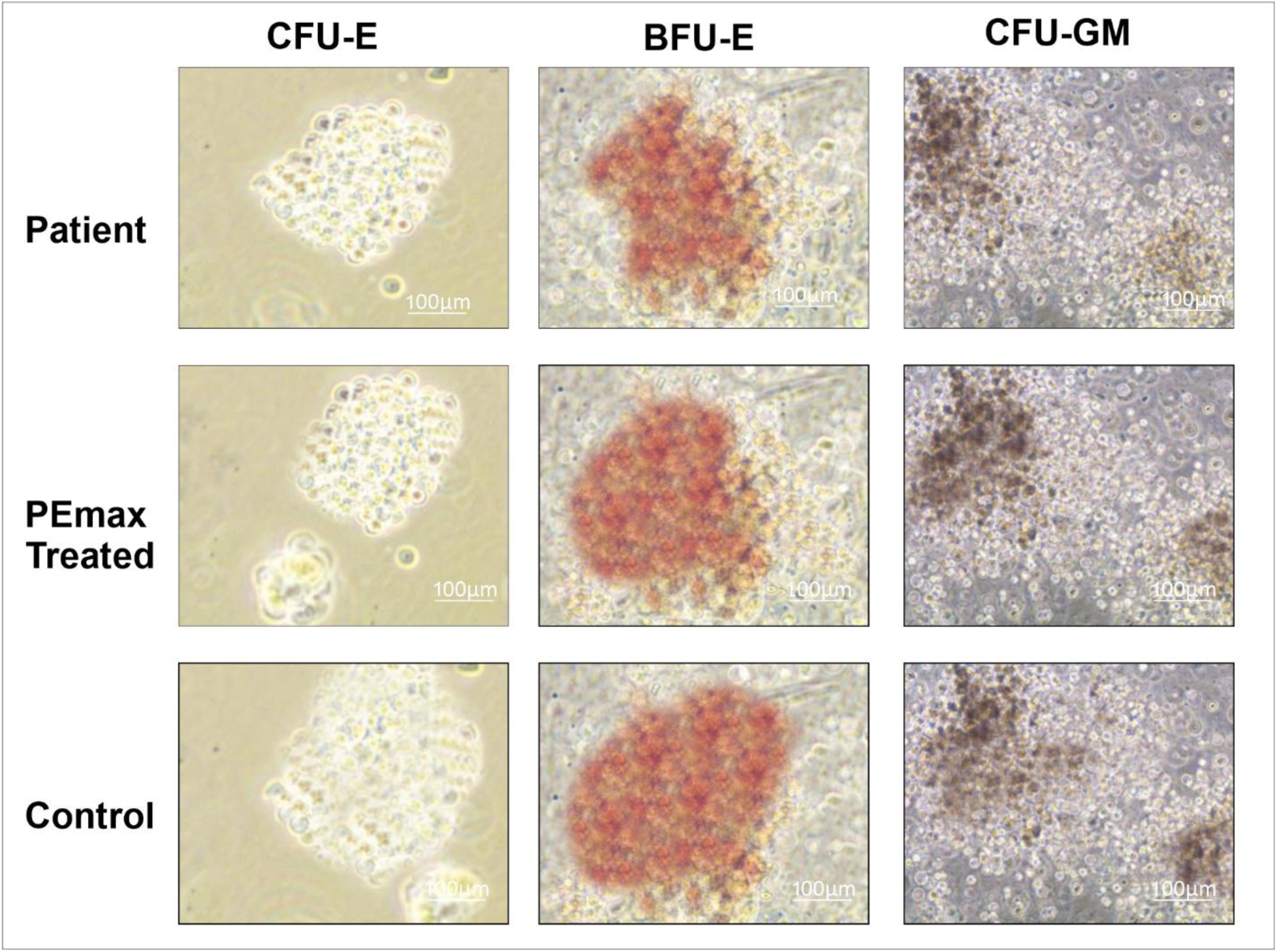
Representative micrographs of haematopoietic colonies (CFU-E, BFU-E, CFU-GM) generated from iPSC-derived haematopoietic progenitors of patient, PEmax-treated, and control lines, illustrating characteristic colony morphology and haemoglobinisation (red, BFU-E) across groups (scale bars, 100 µm).

**Supplementary Figure S2.**
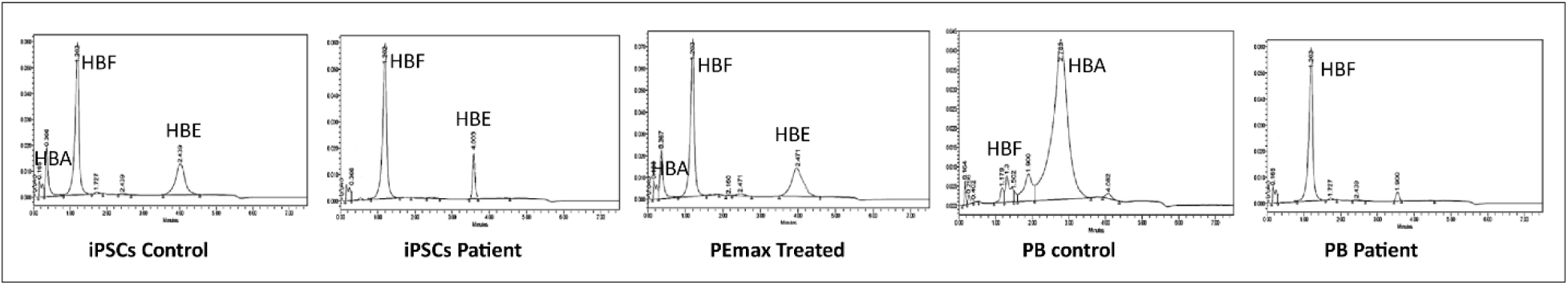
HPLC chromatograms of haemoglobin fractions. Representative HPLC traces showing haemoglobin peak profiles (HBA, HBF, HBE) in iPSC-derived erythroid cells (control, patient, PEmax-treated) and peripheral blood samples (control, patient), corresponding to the quantified haemoglobin fractions shown in the main figure.

**Supplementary Table S1.**
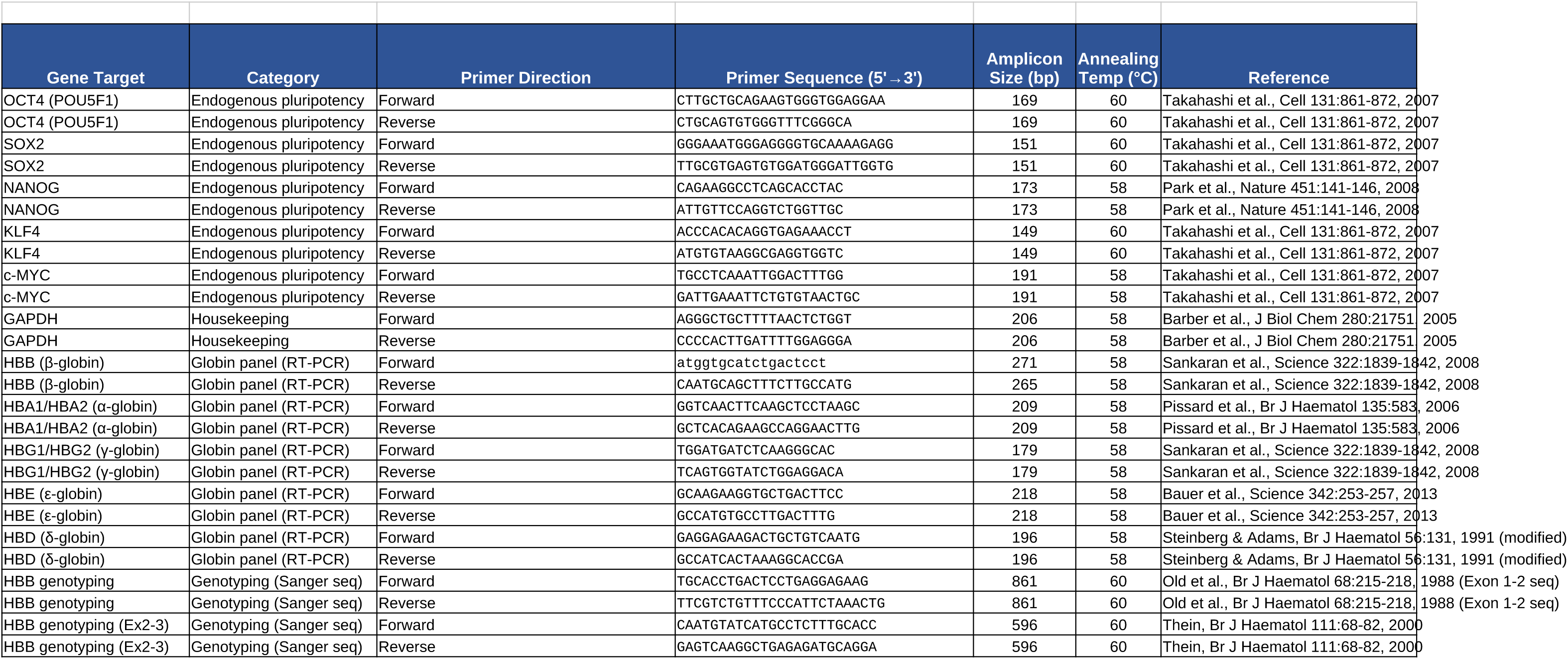
Primer Sequences for RT-PCR and qRT-PCR.

**Supplementary Table S3.**
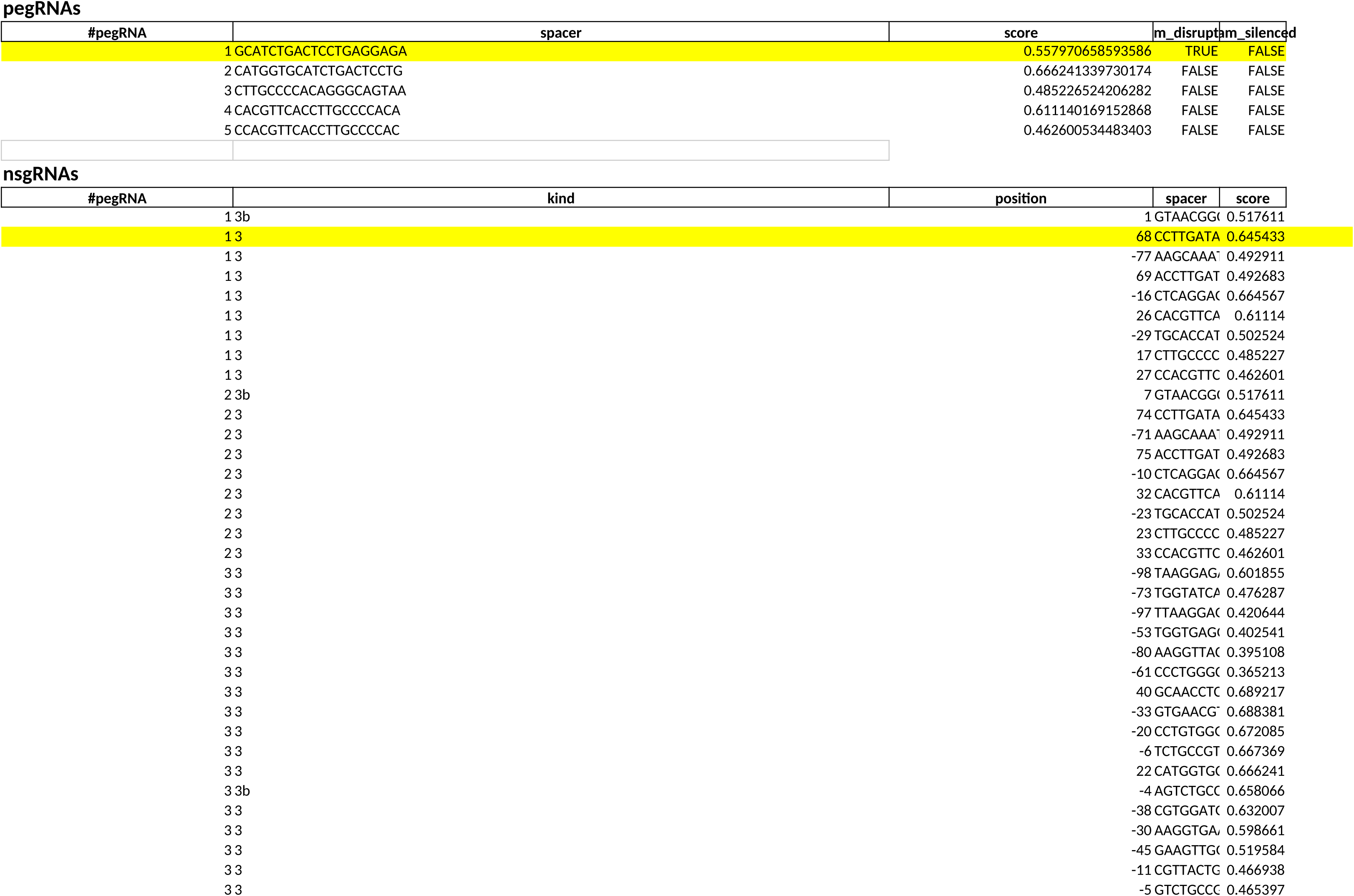
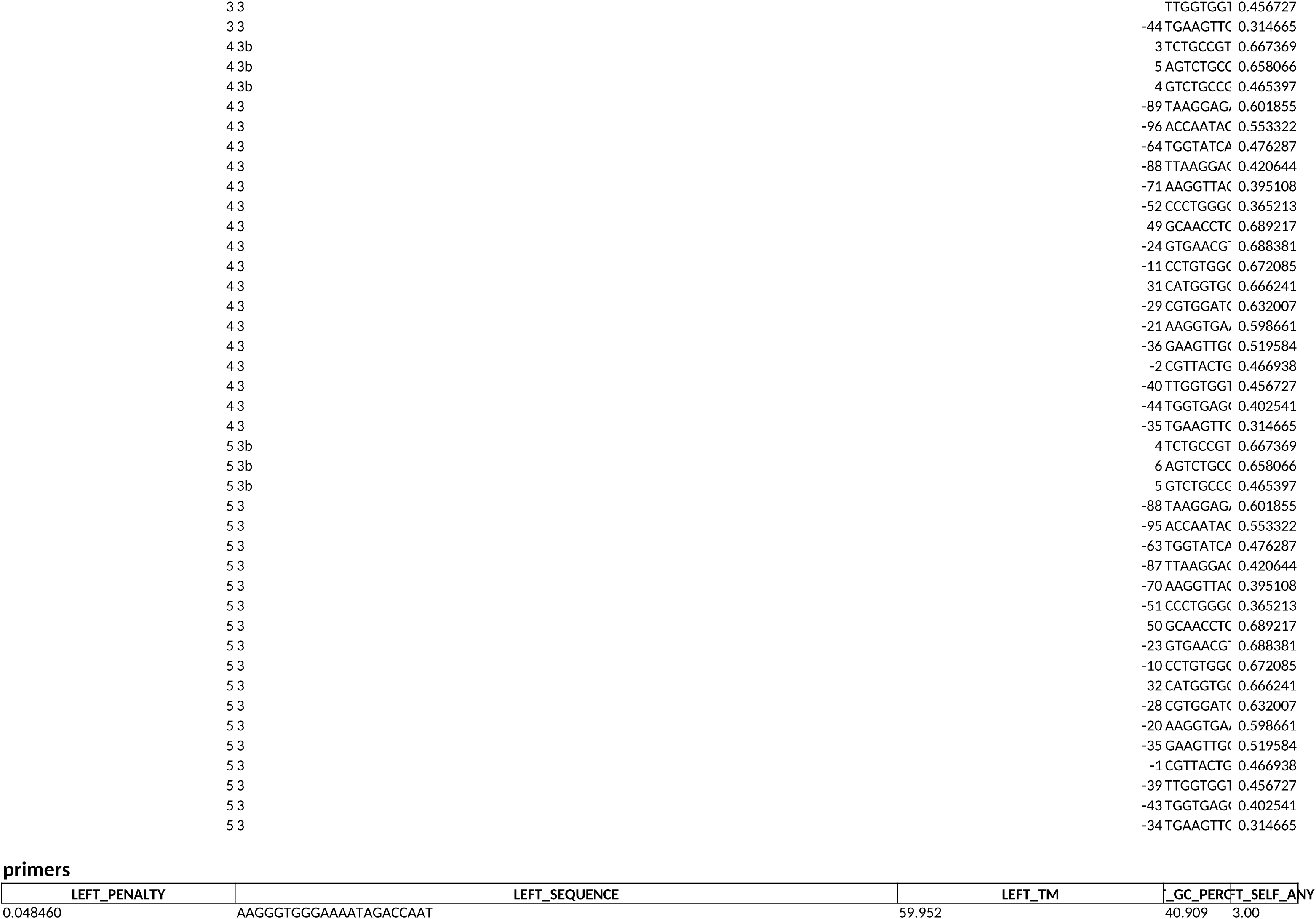
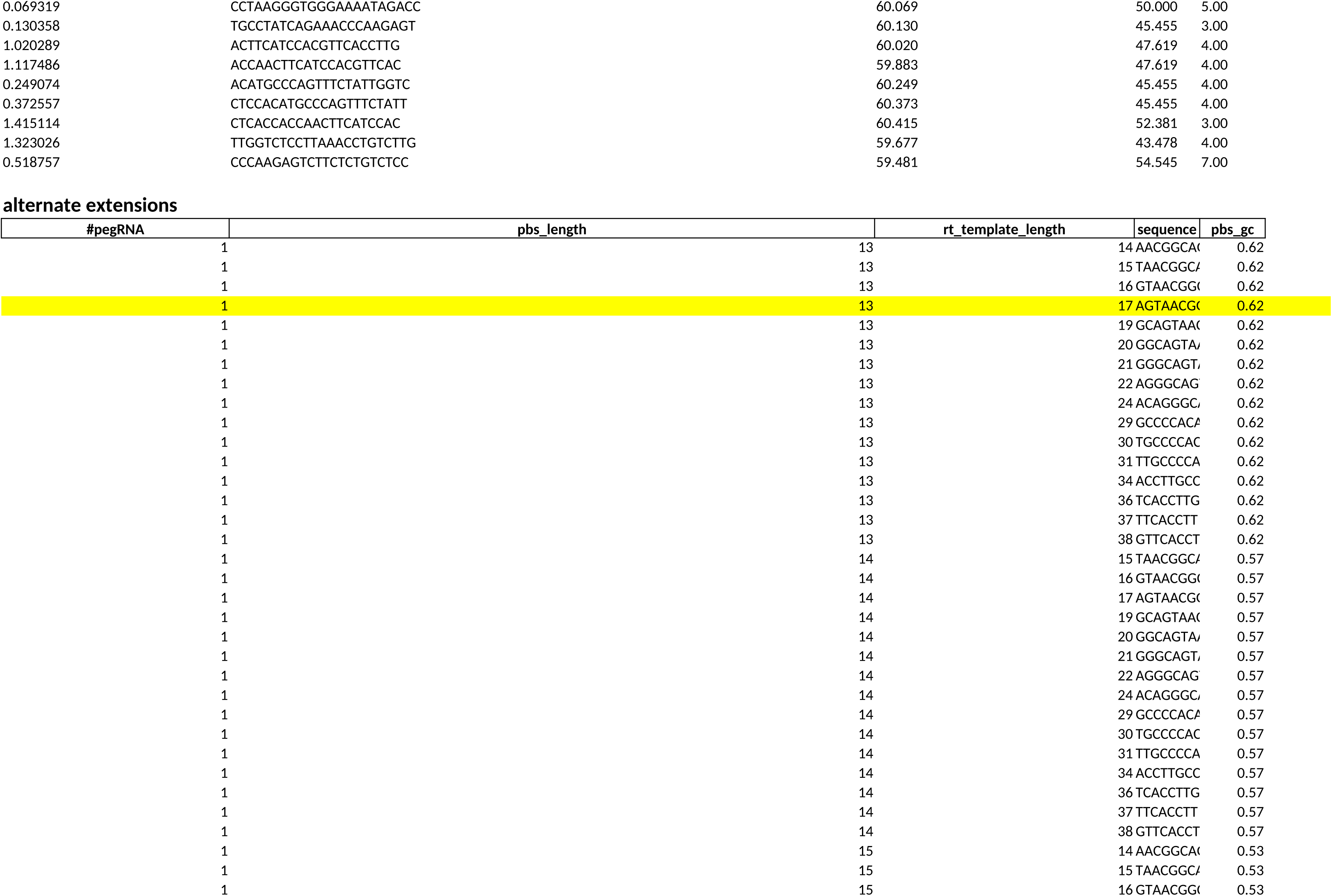
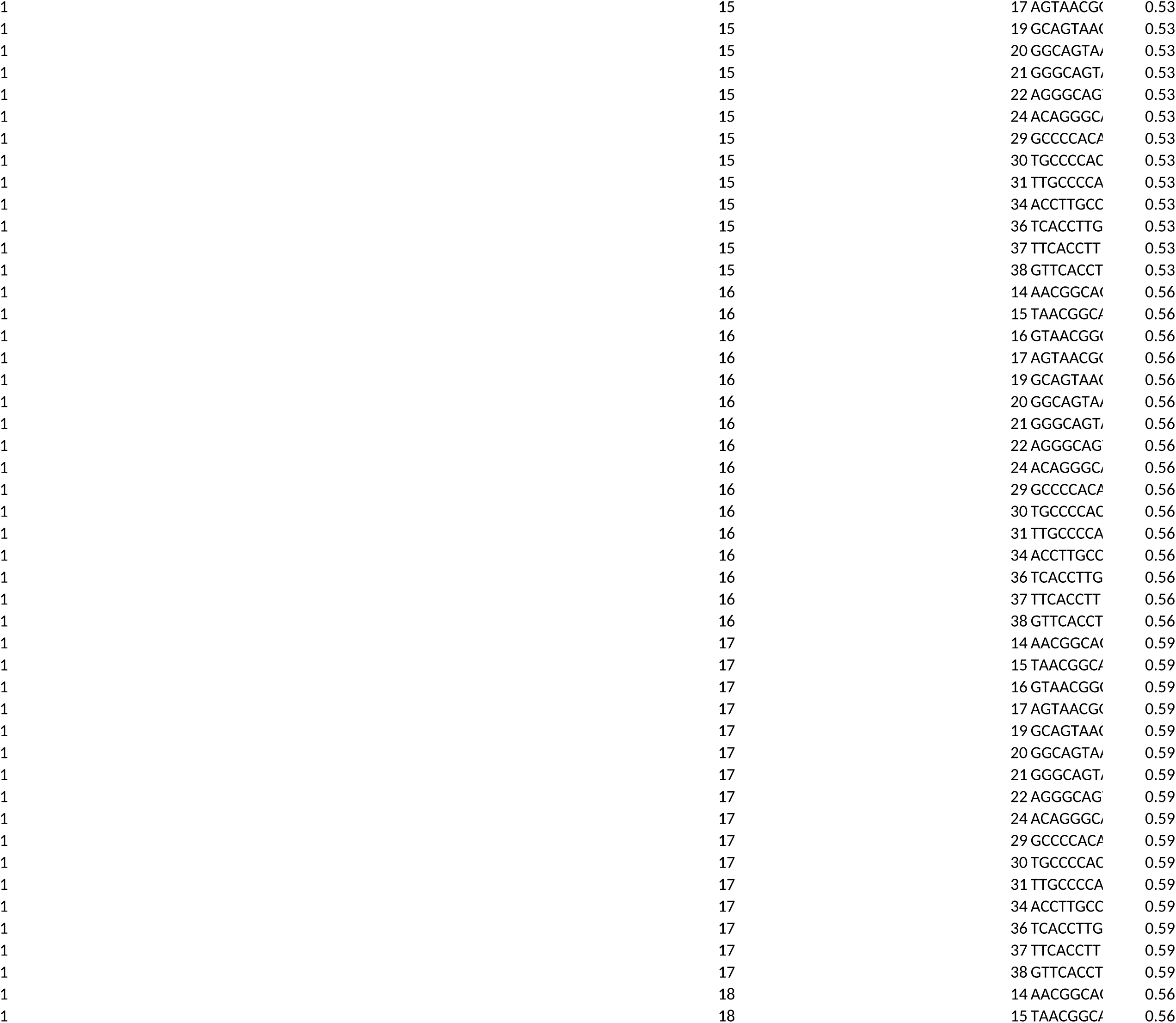
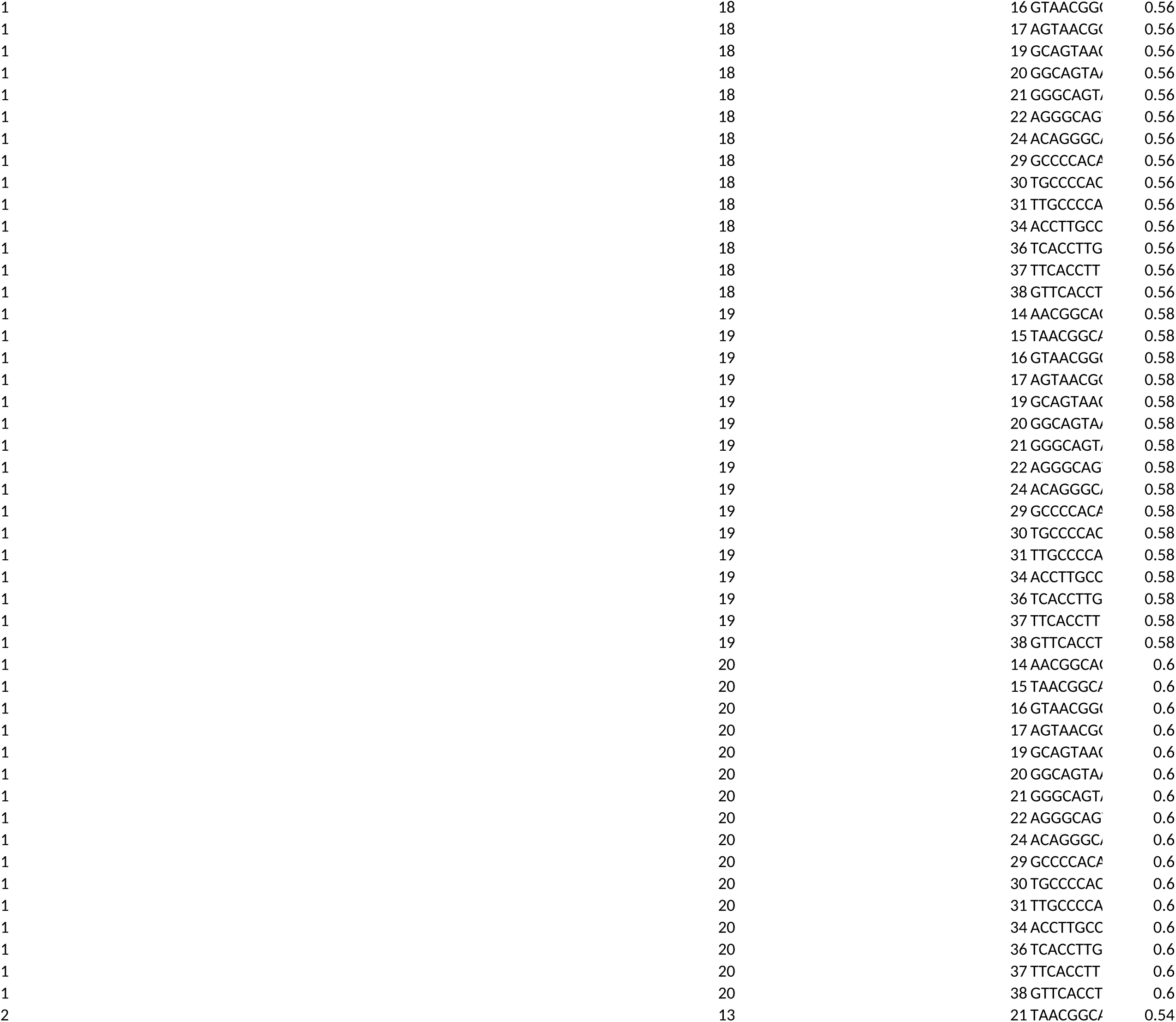
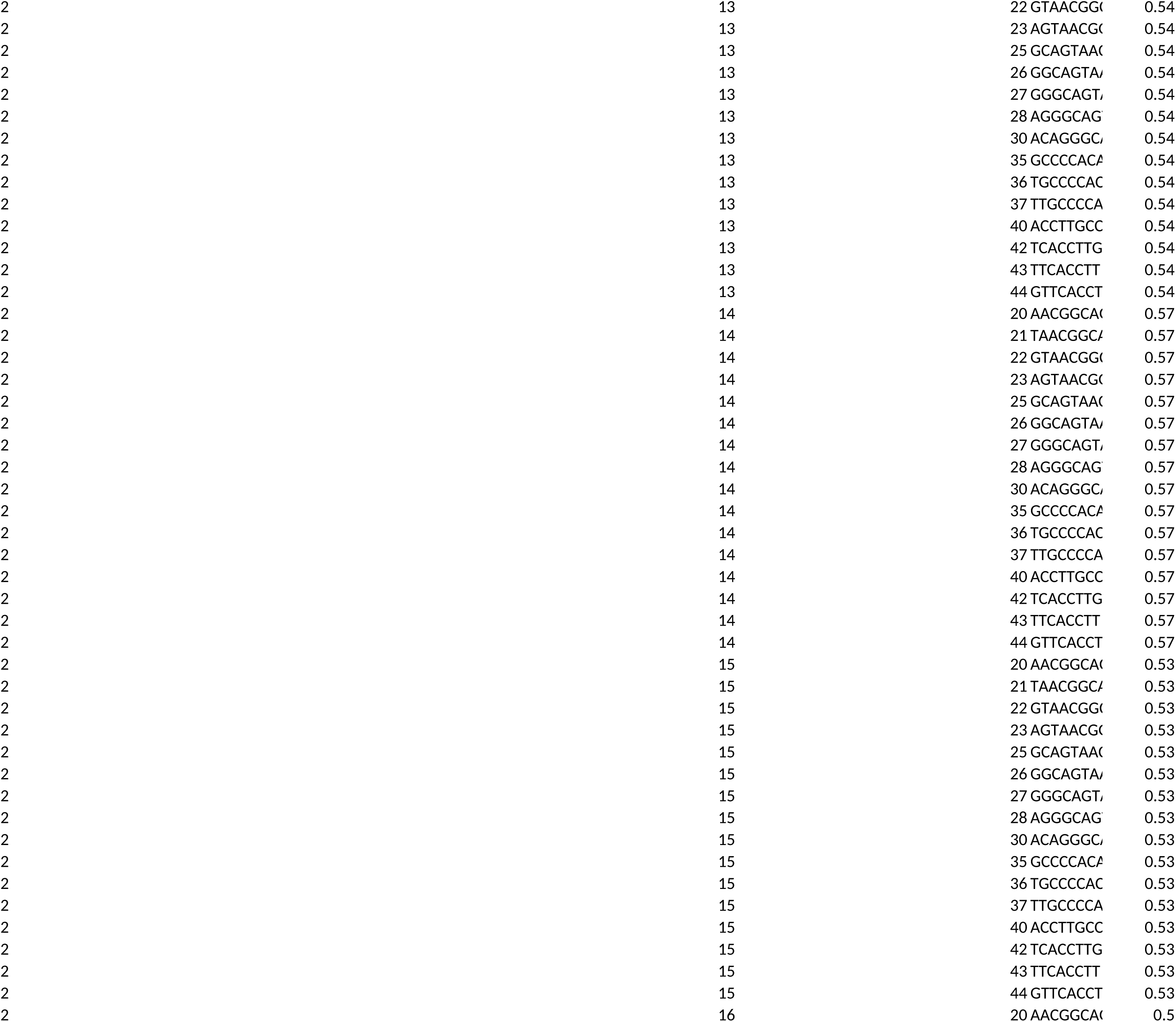
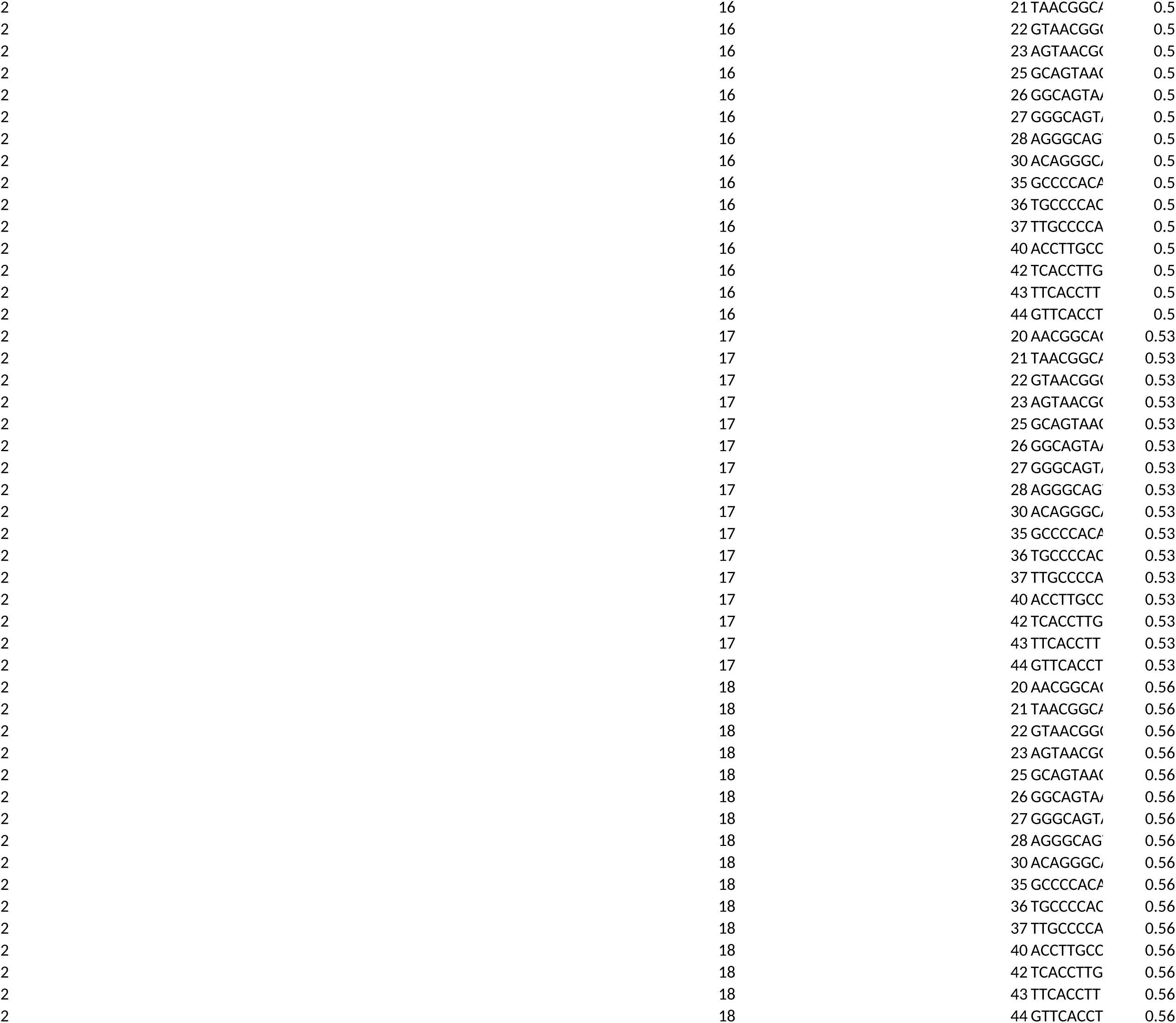
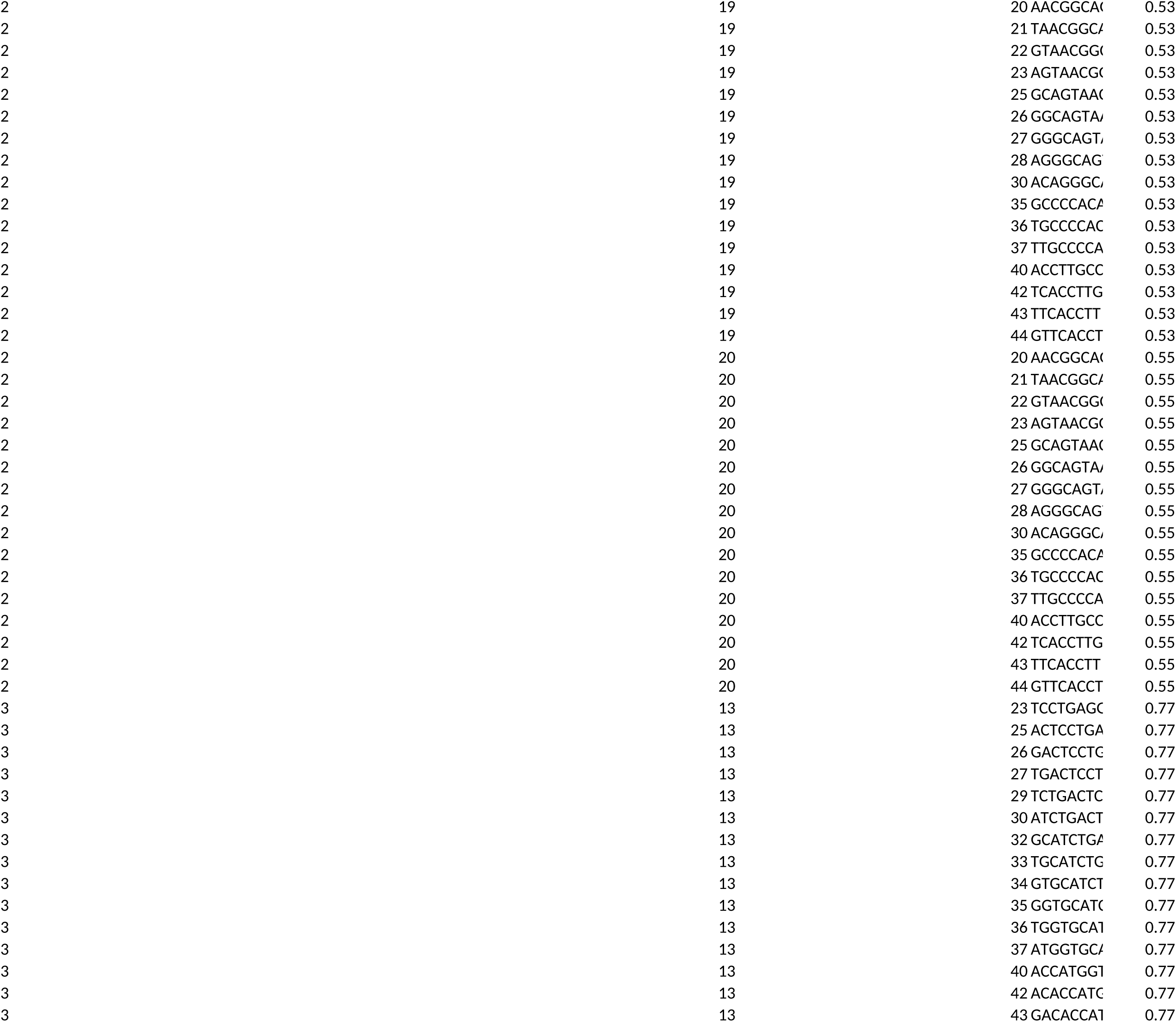
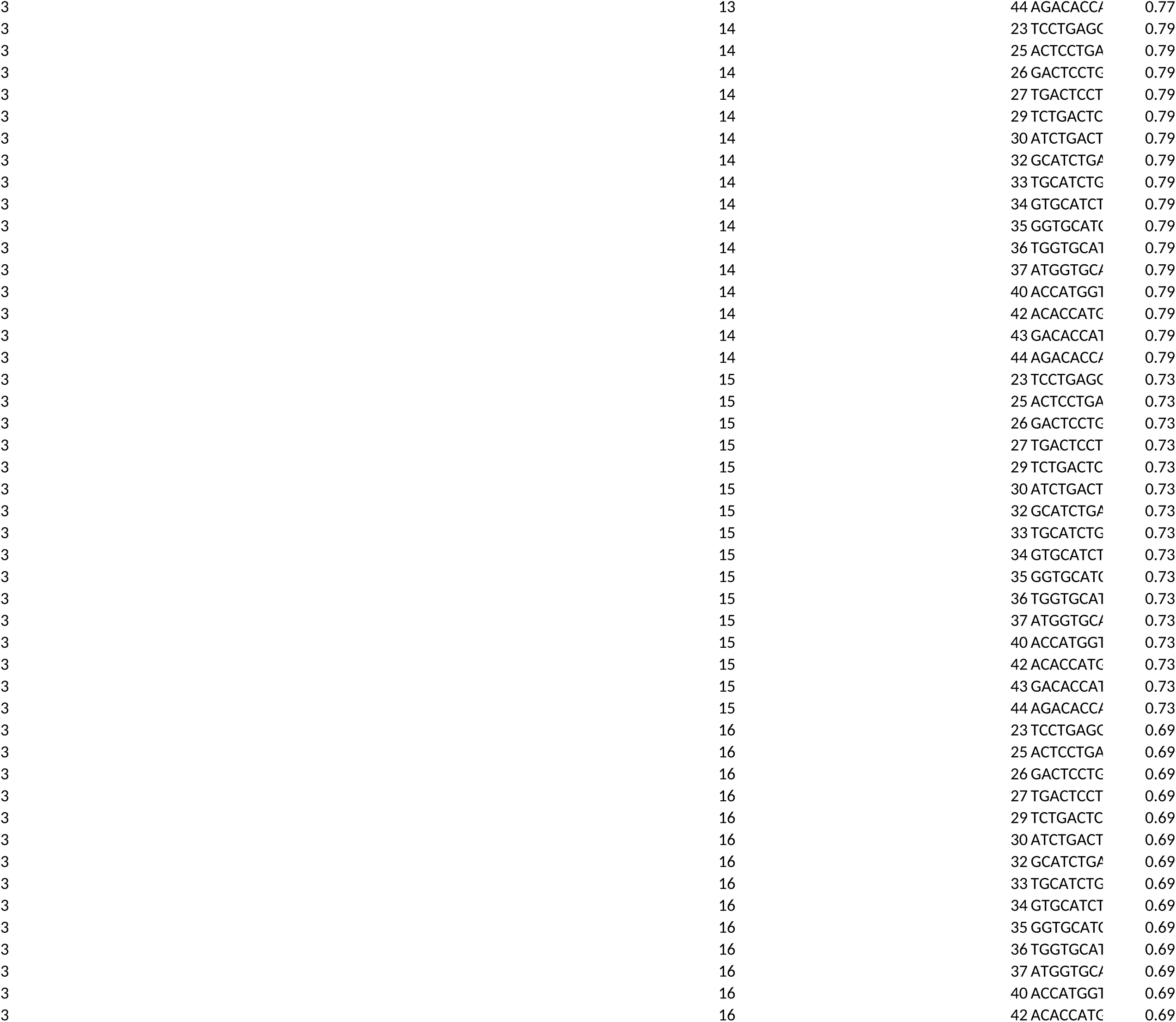
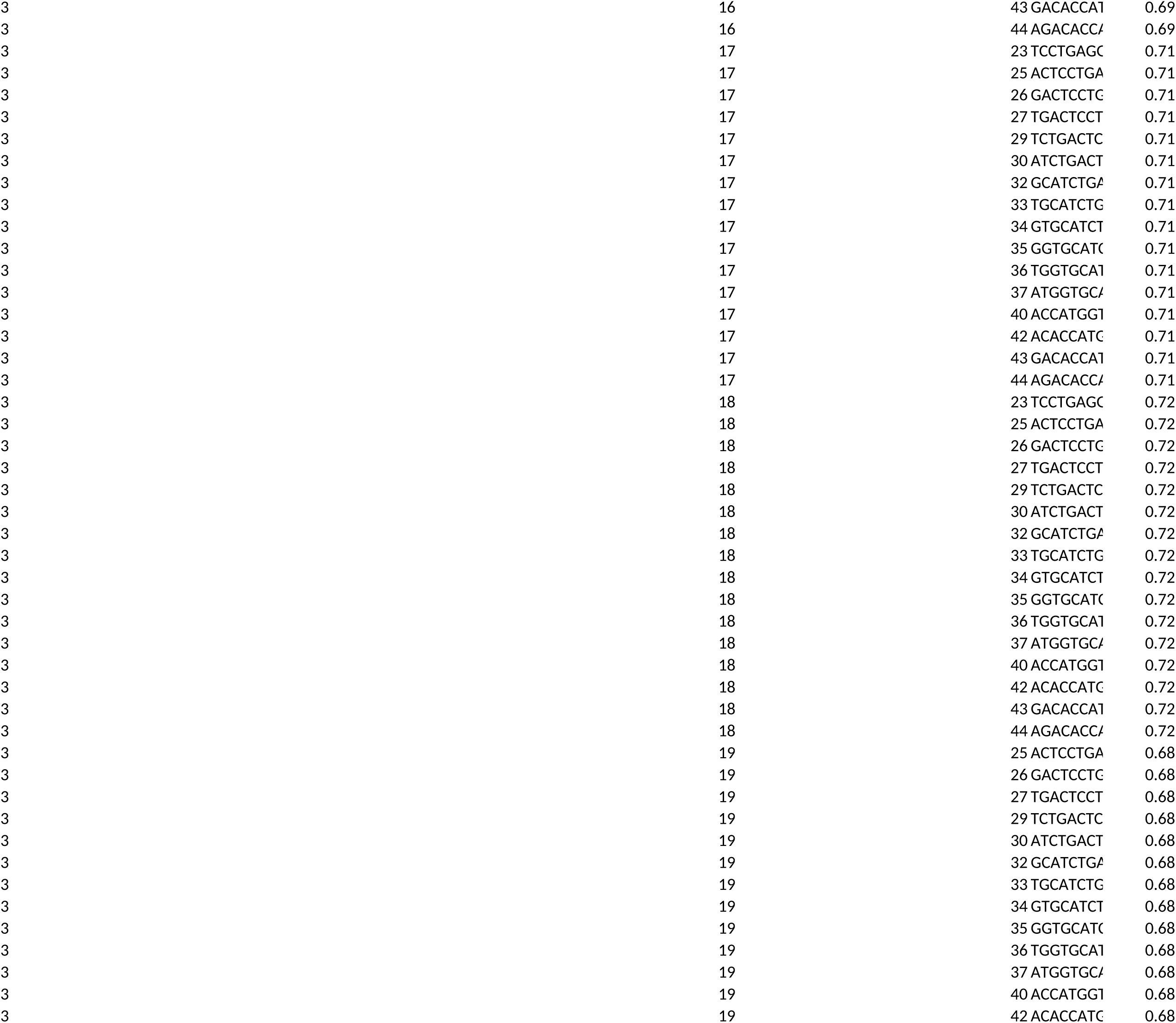
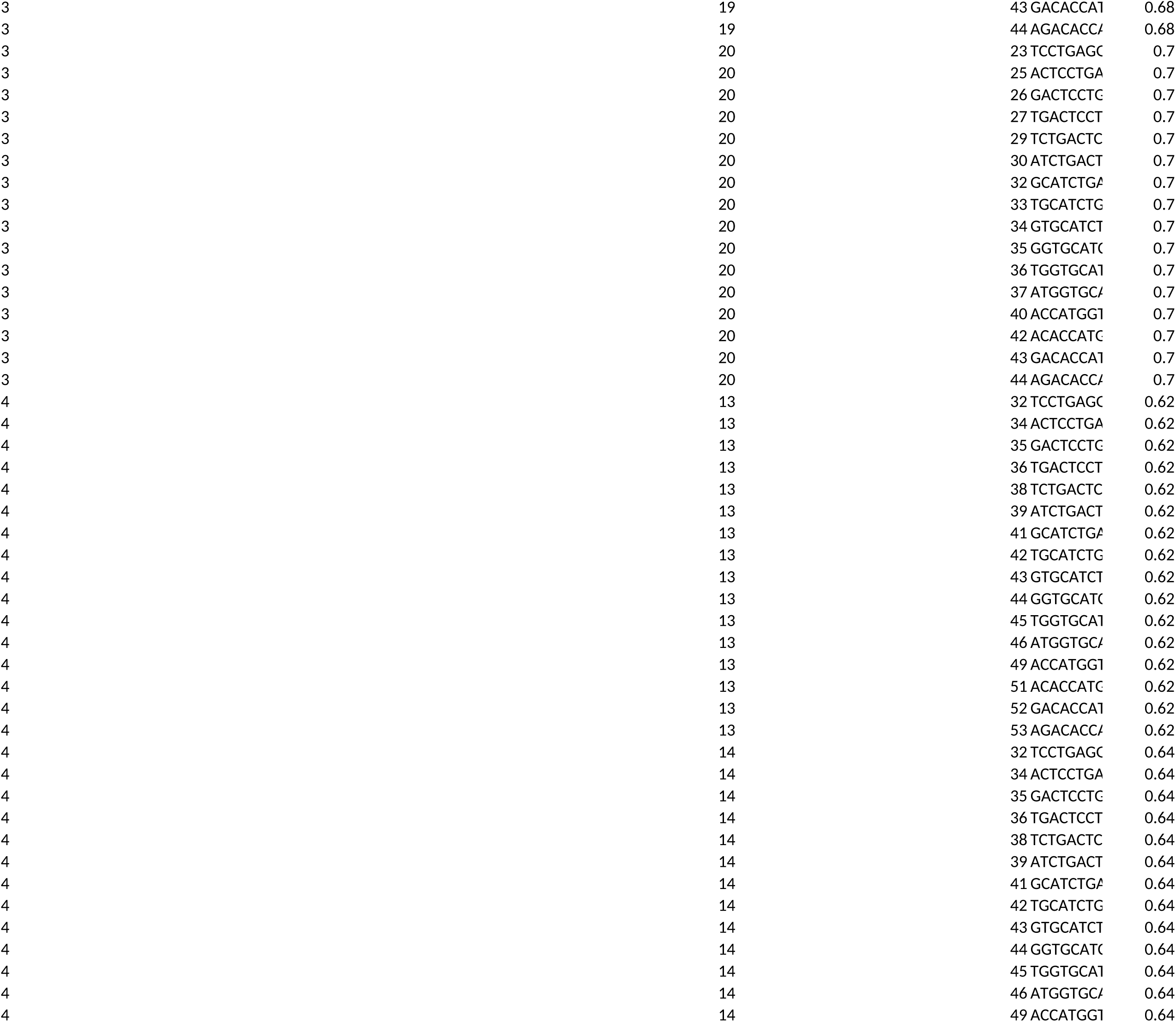
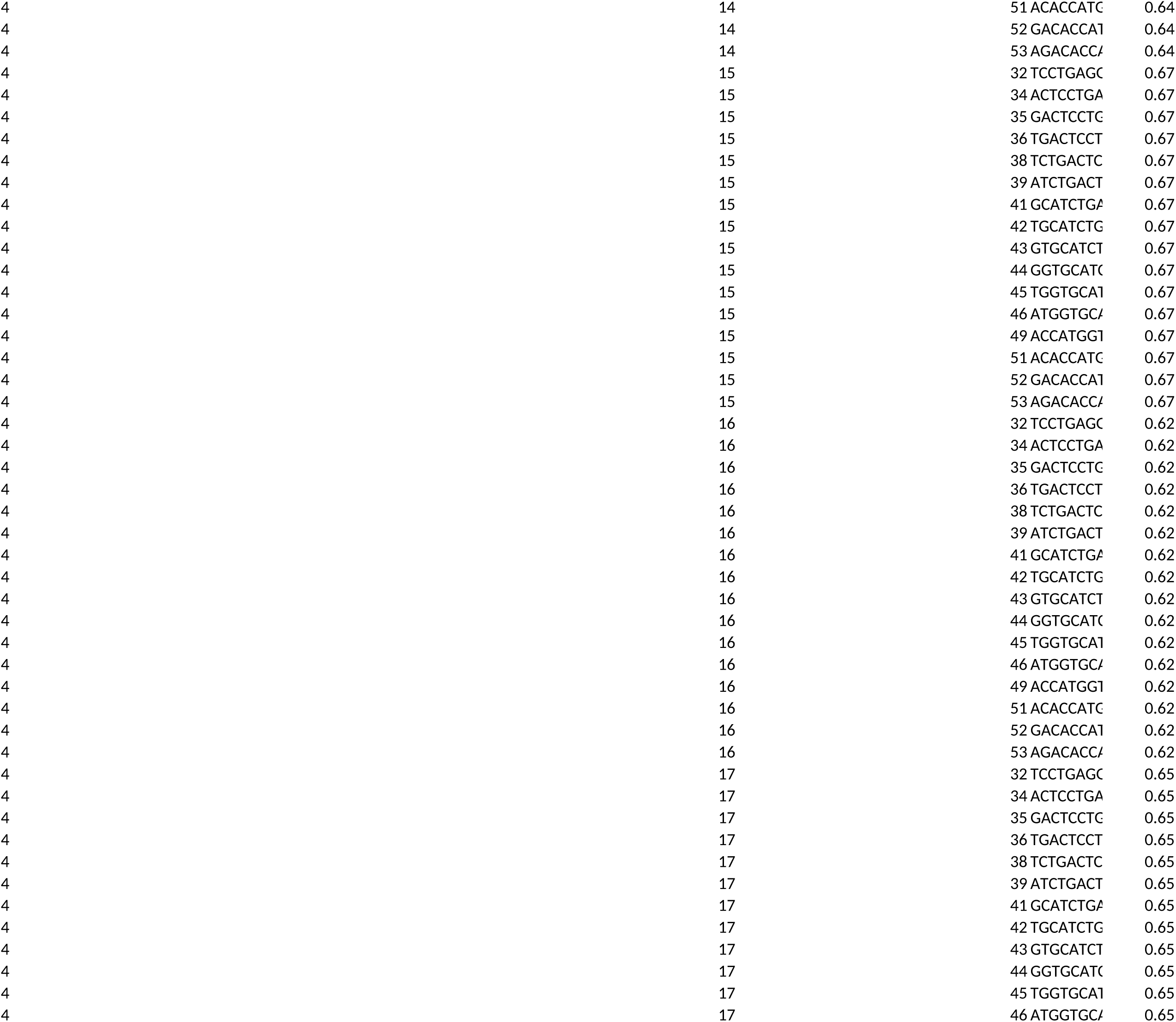
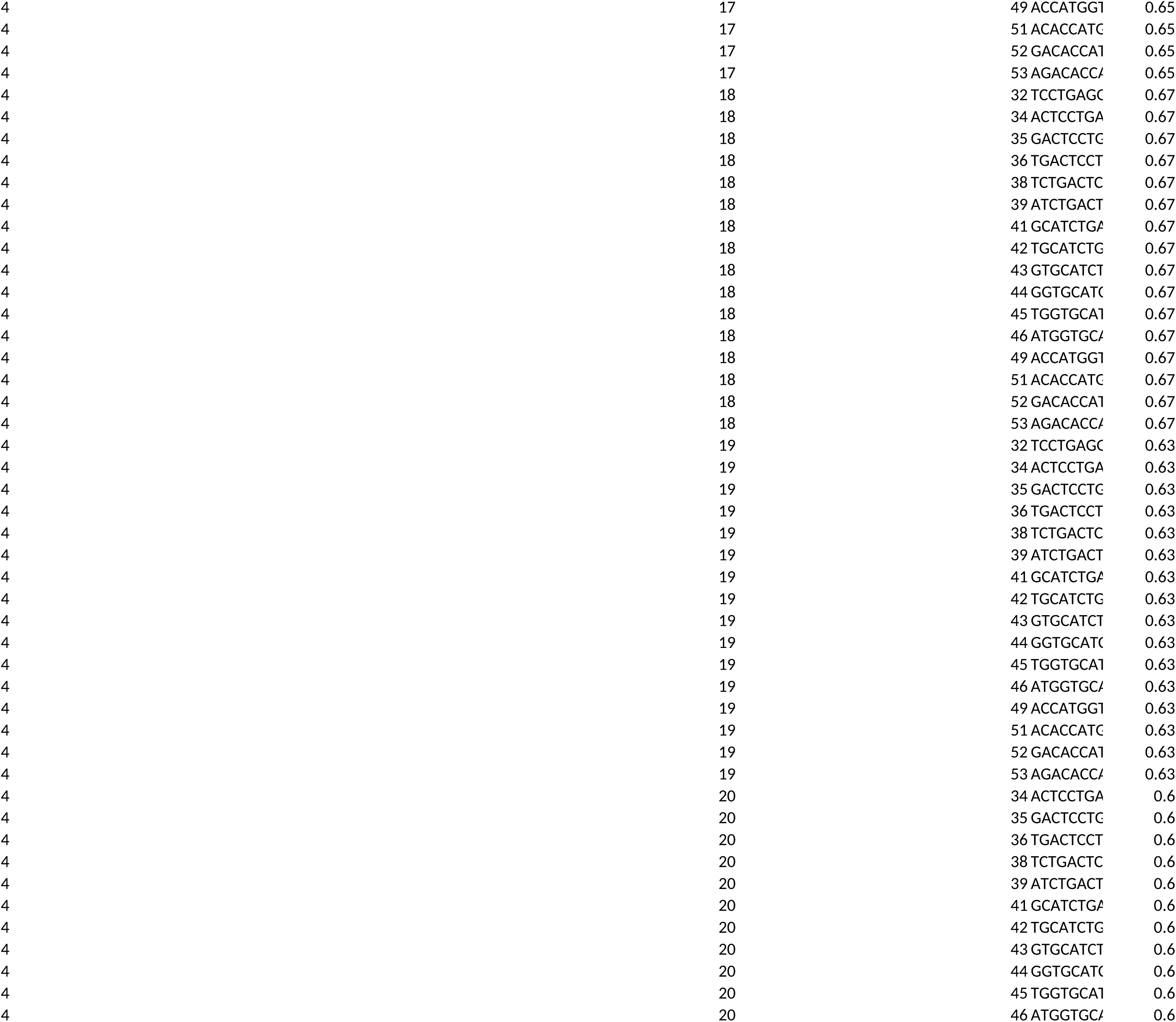
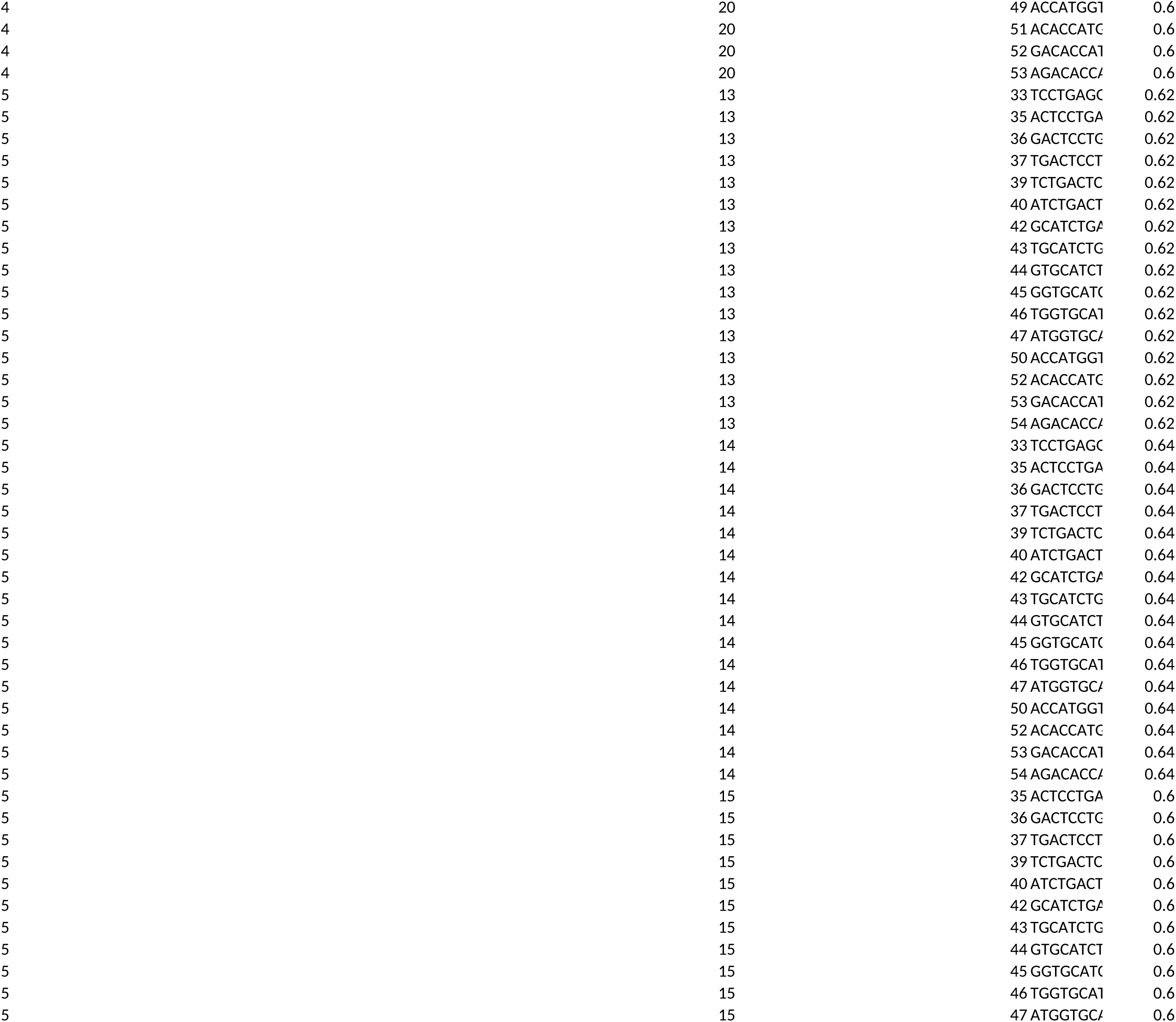
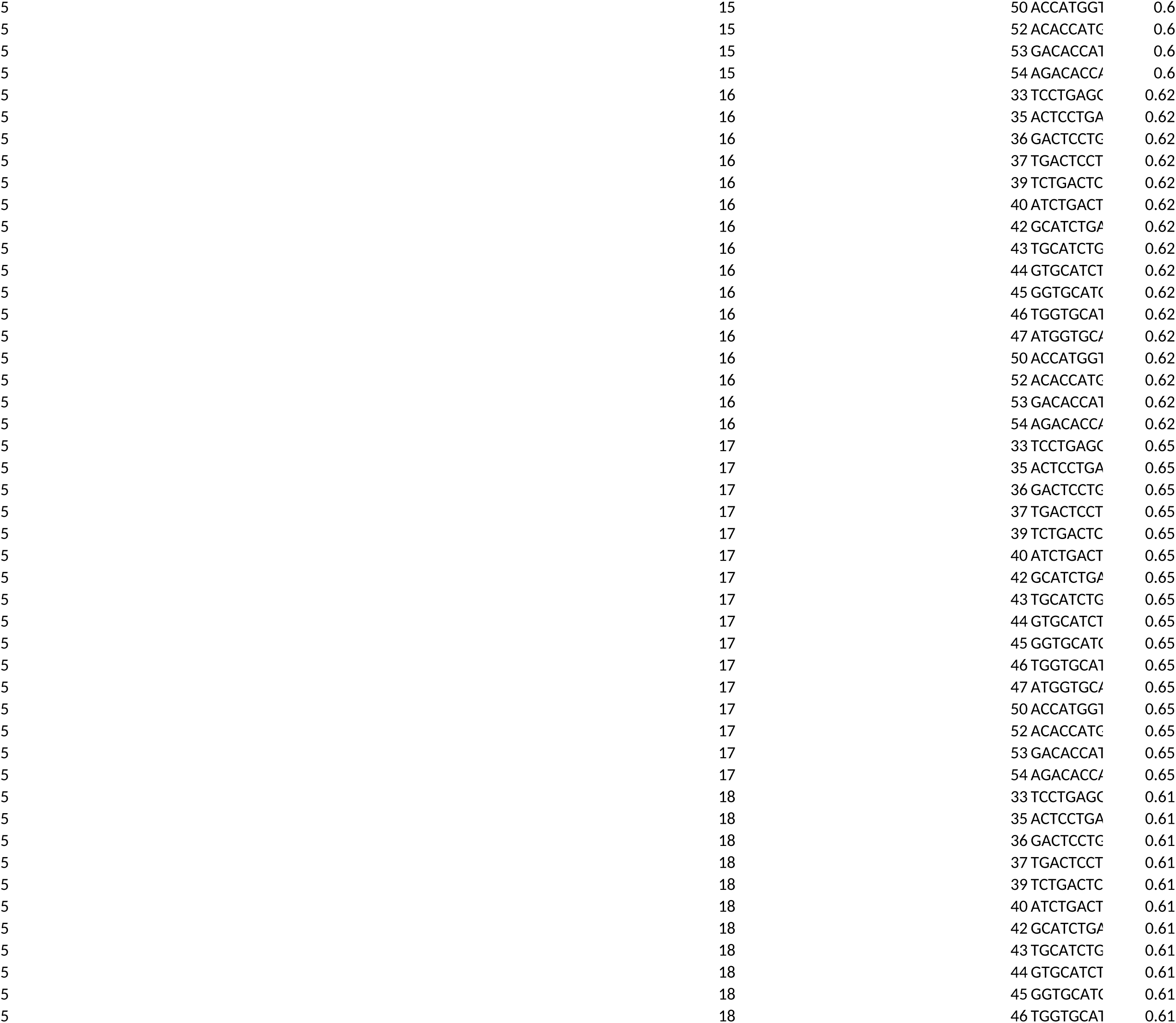
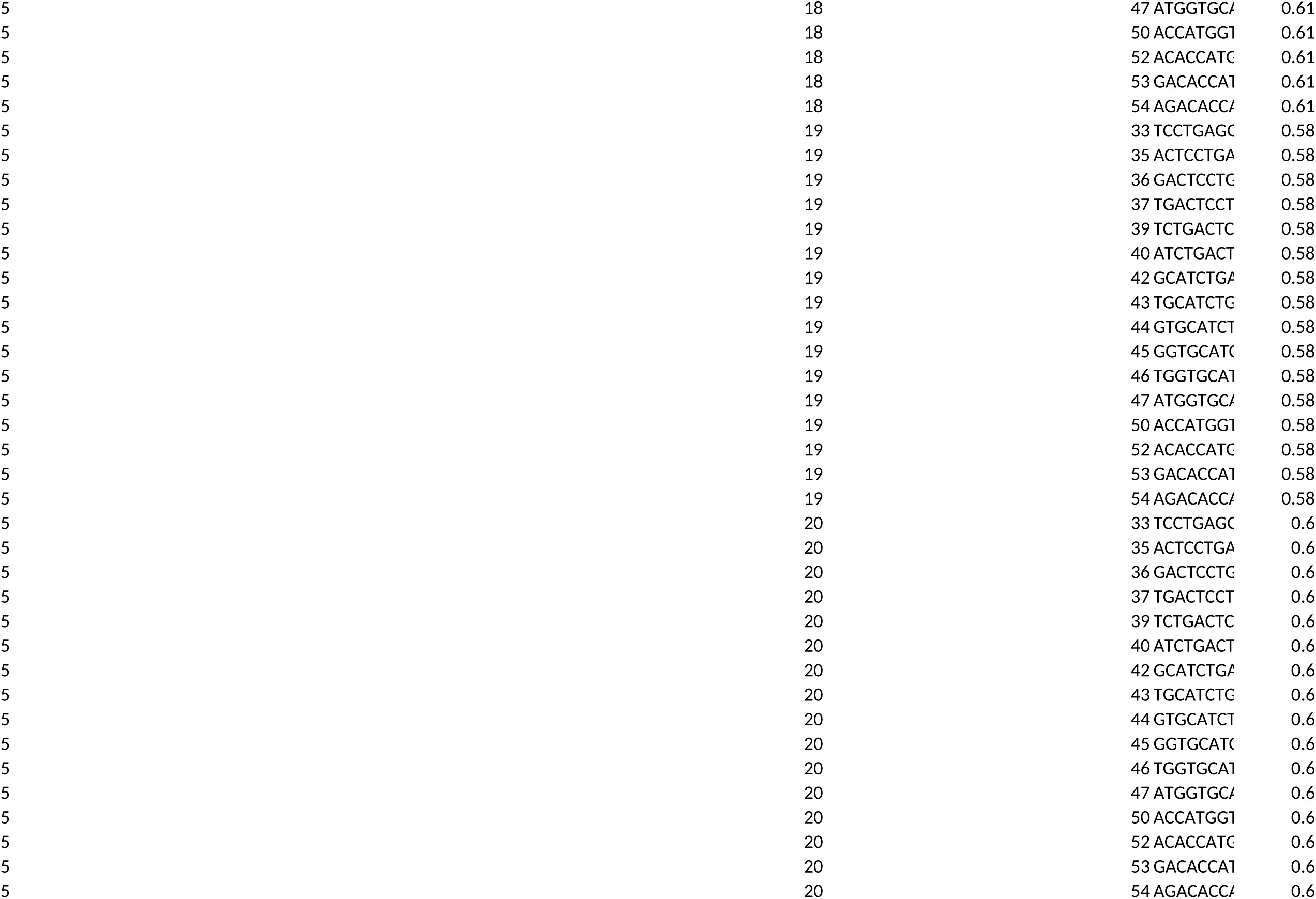
Prime Editing Guide RNA (pegRNA) and Nicking sgRNA Design.

**Supplementary Table S4.**
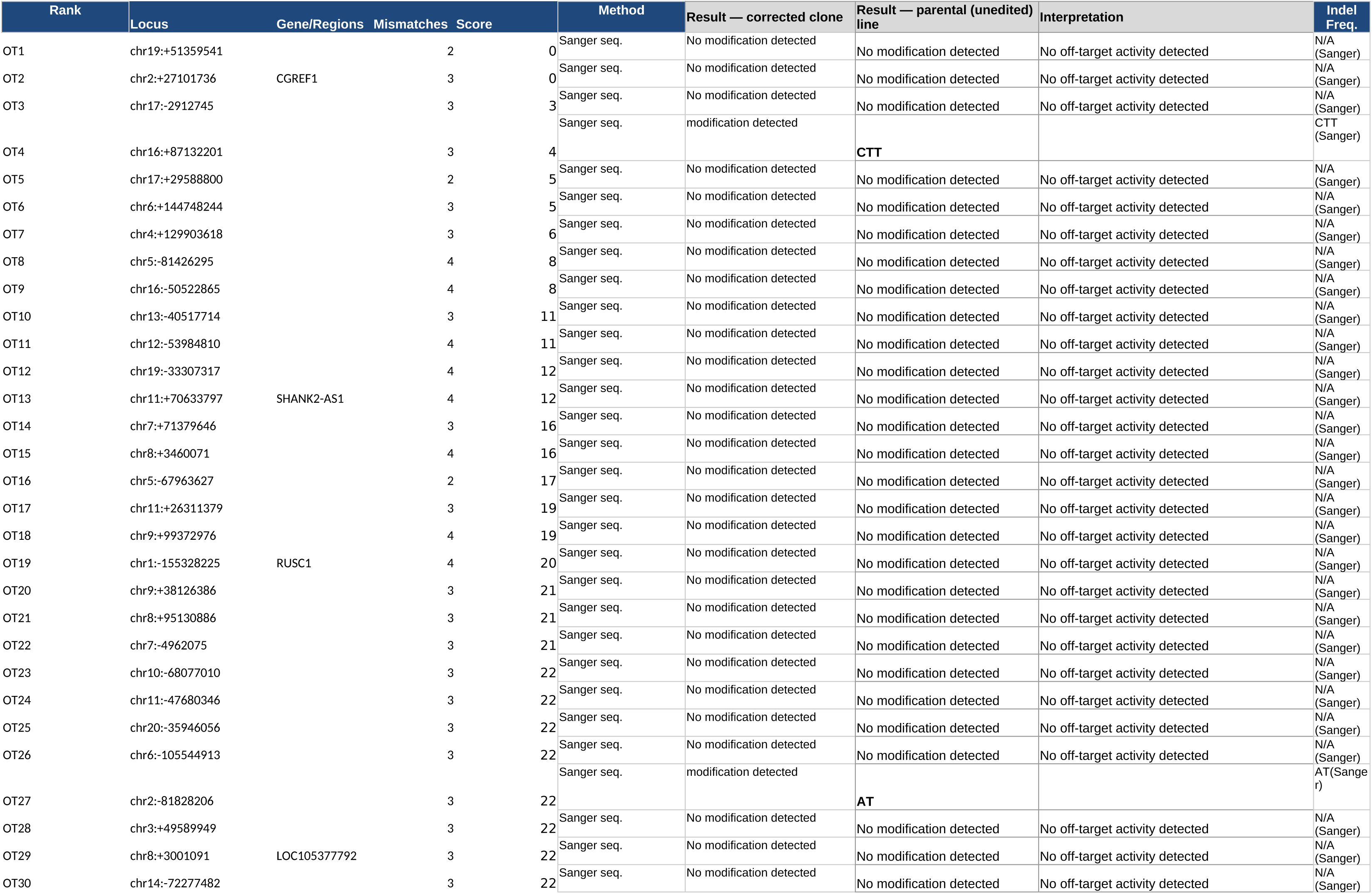
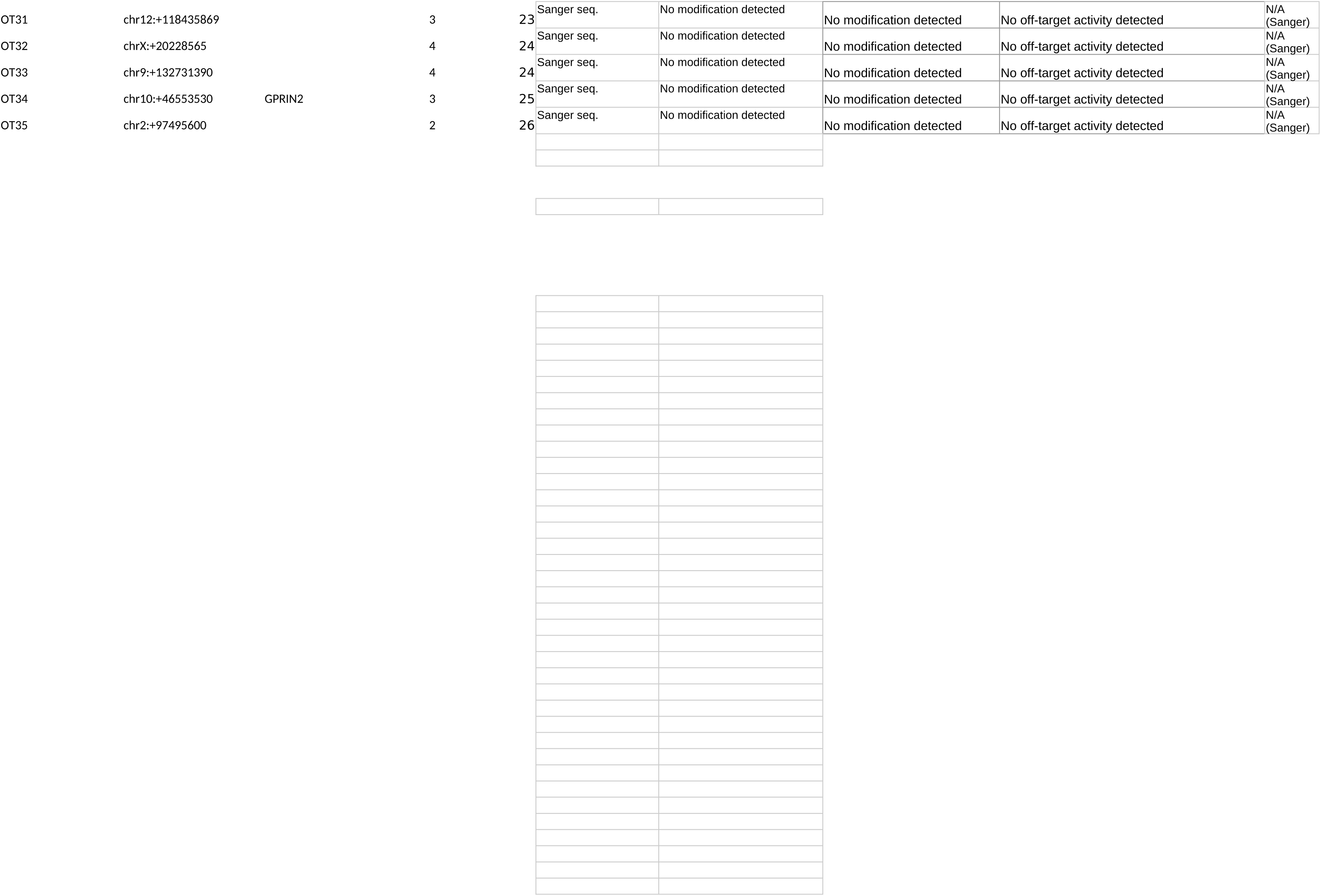
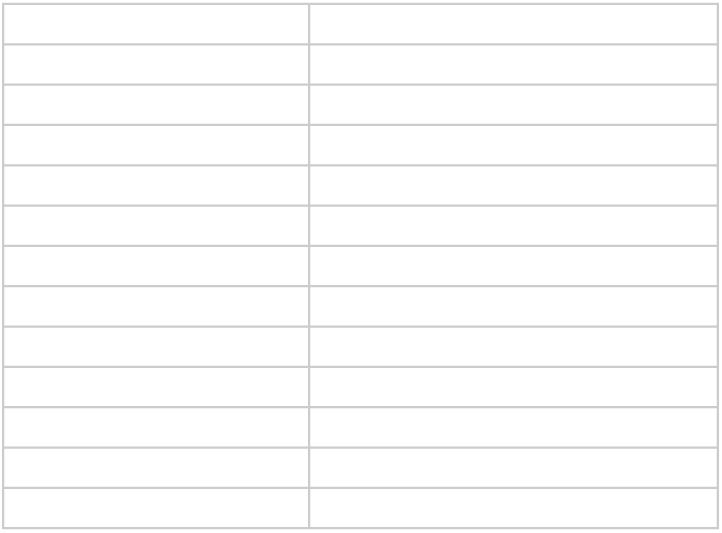
Predicted Off-Target Sites and Sequencing Results.

**Supplementary Table S5.**
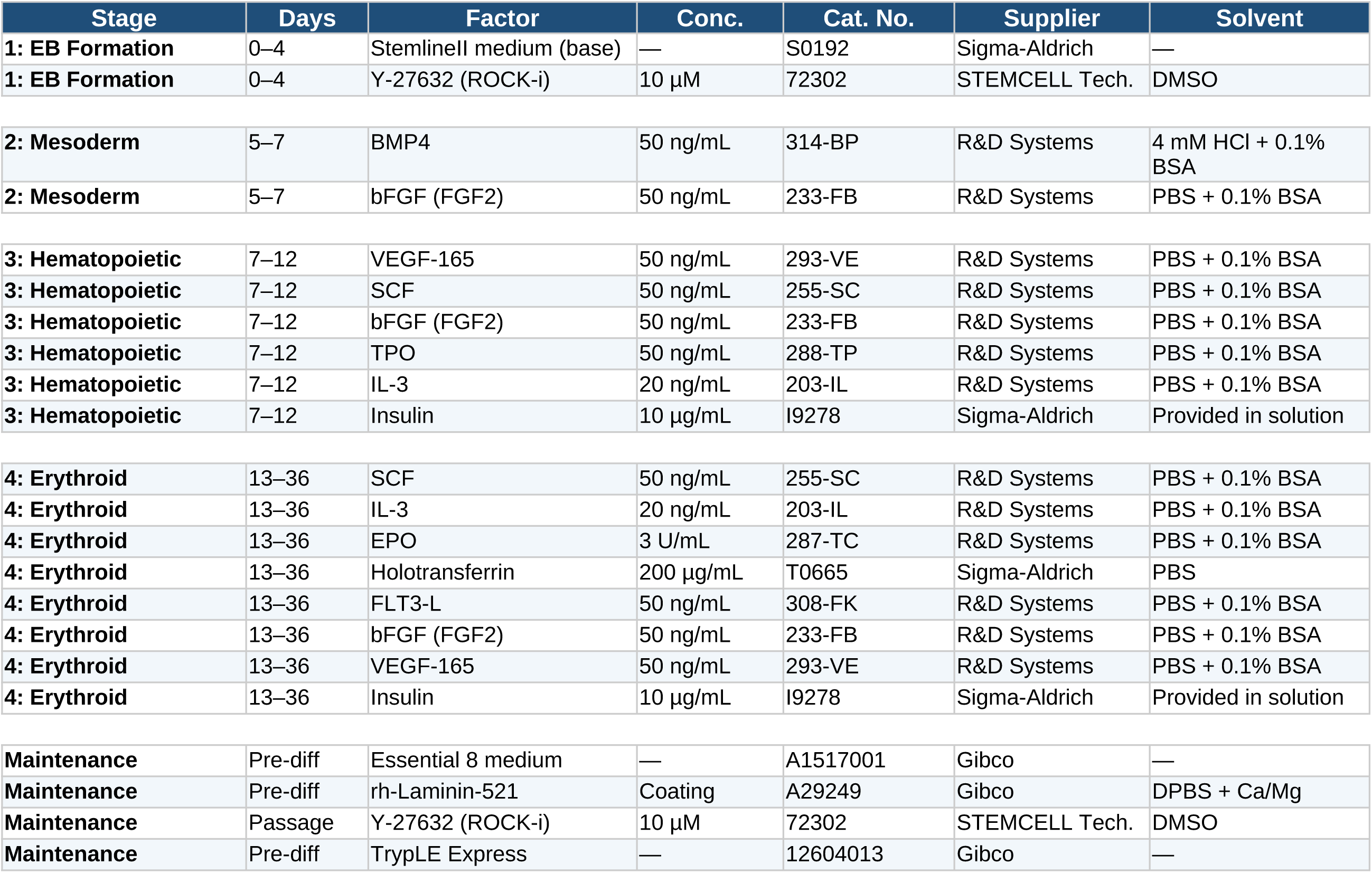
Cytokines, Growth Factors, and Media for Hematopoietic Differentiation.

**Supplementary Table S6.**
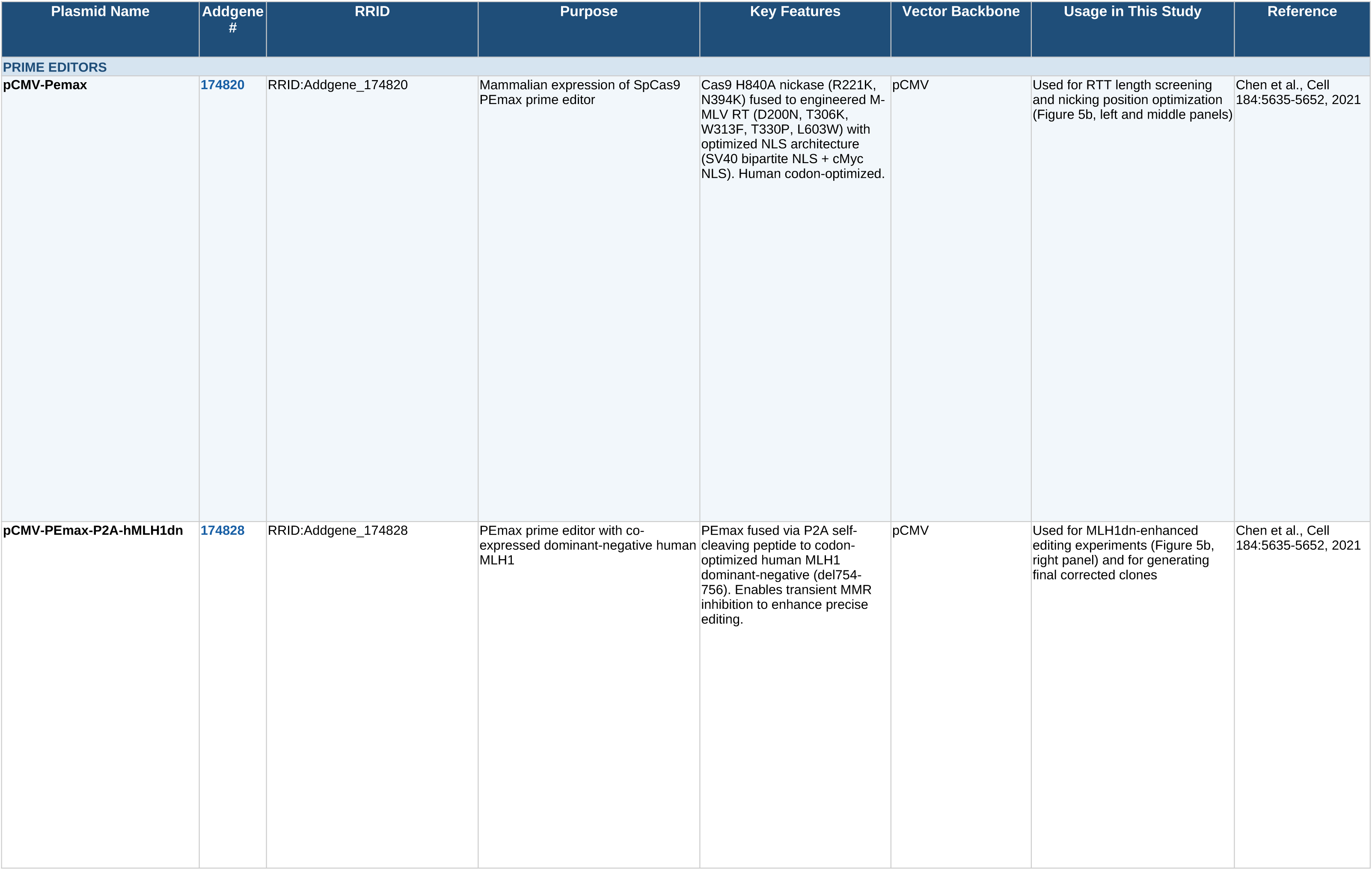
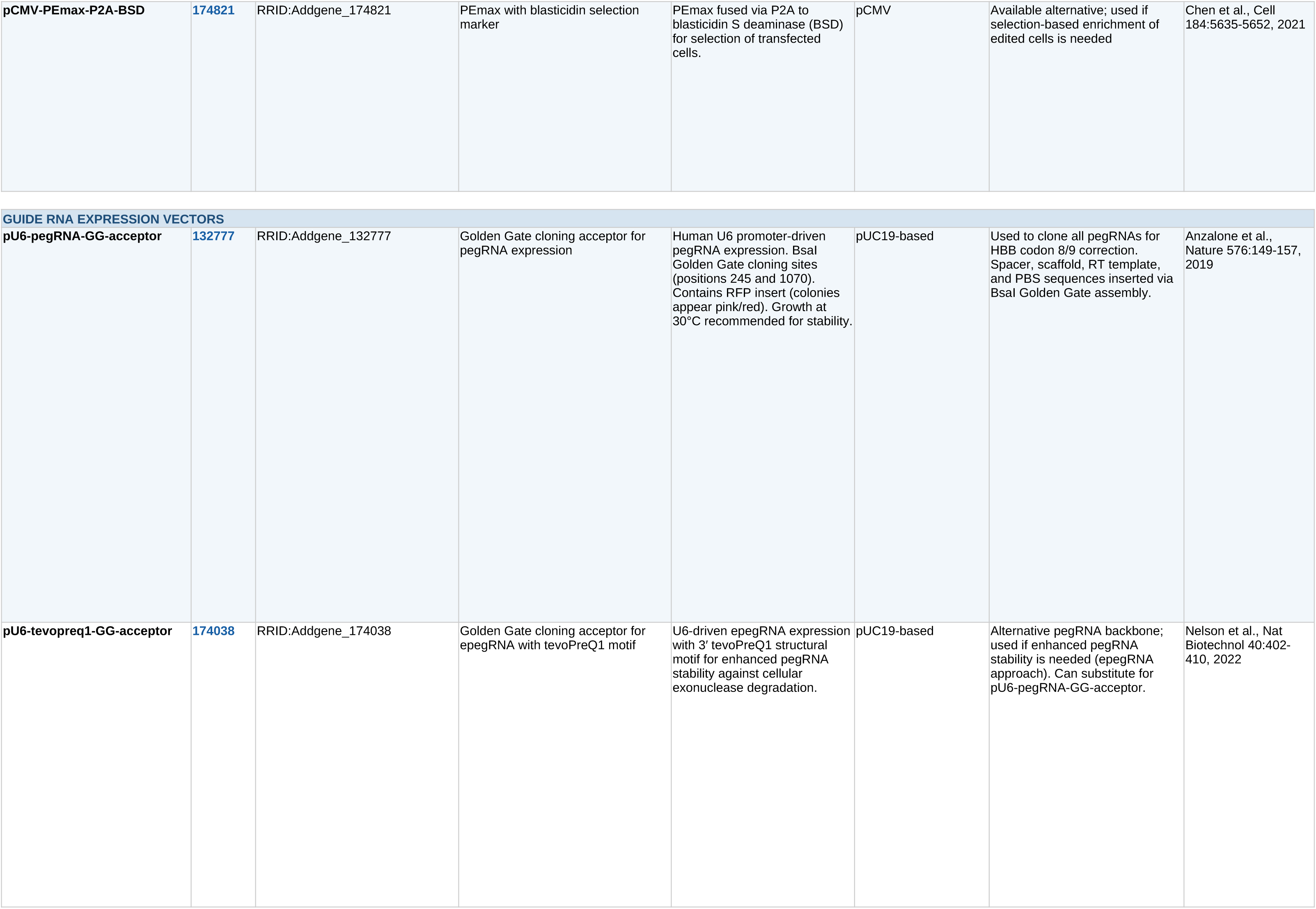
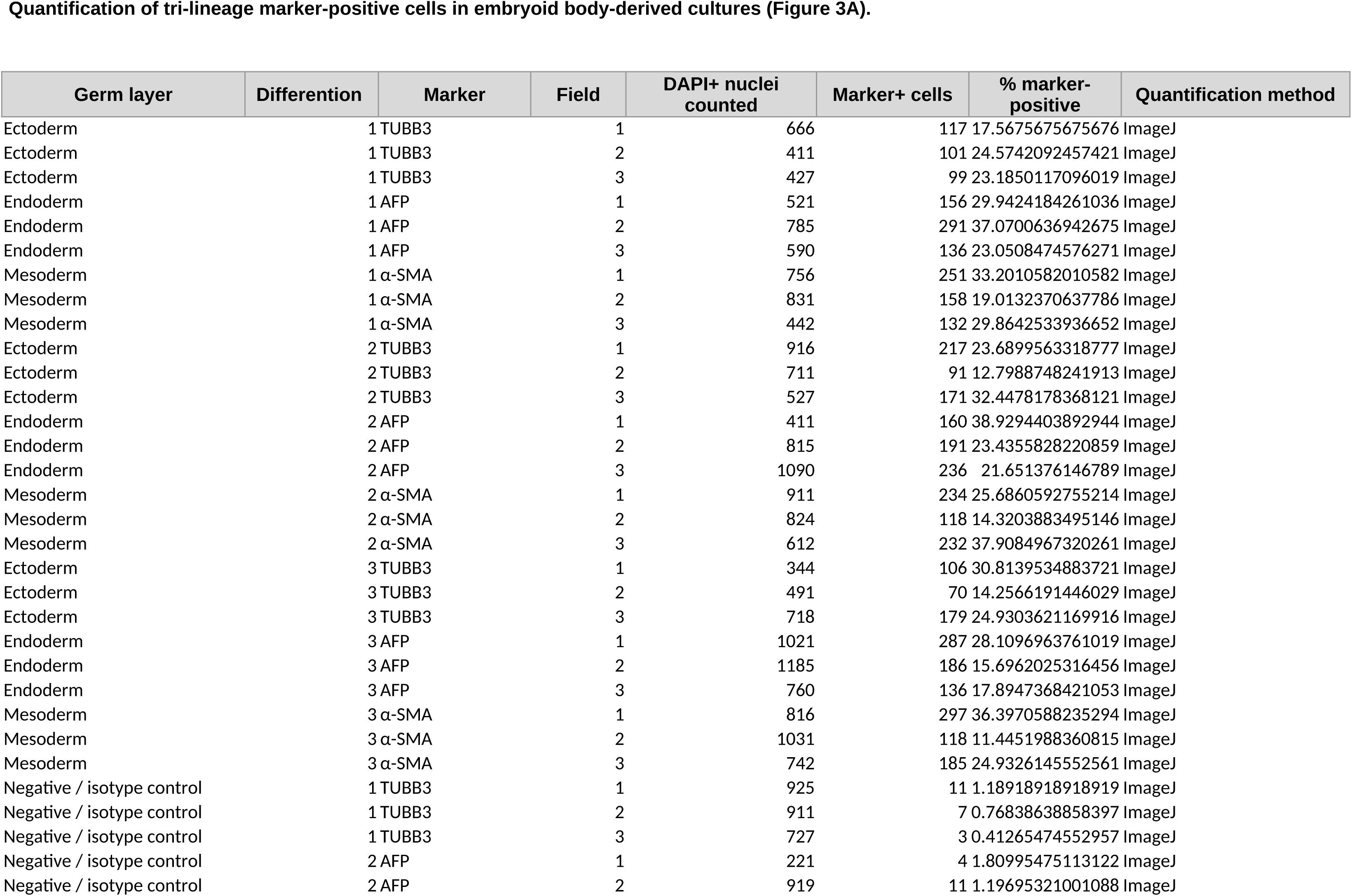
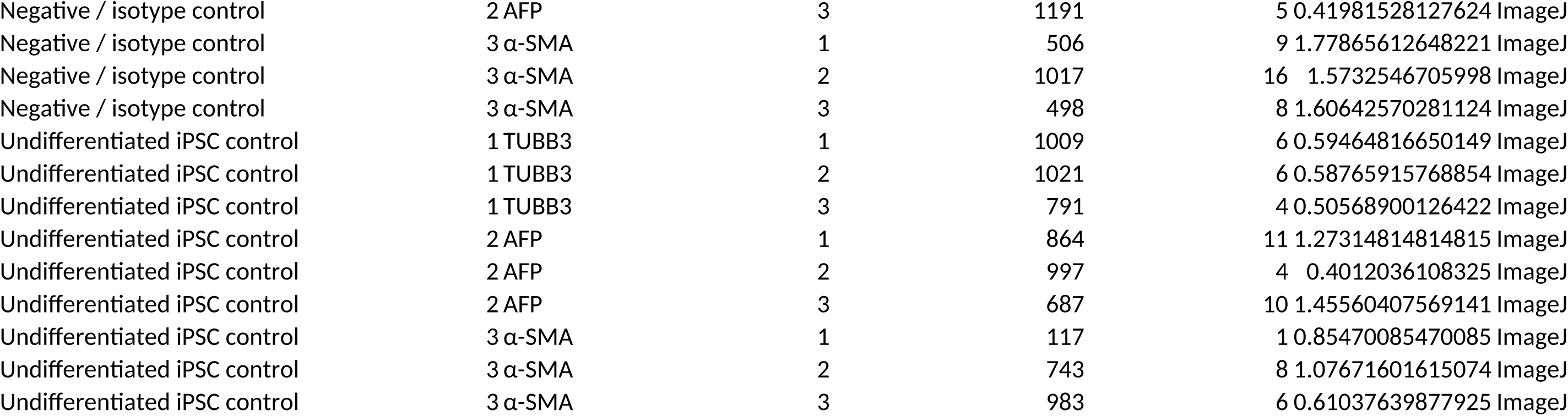
Plasmids Used in This Study. All prime editing plasmids were obtained from Addgene (deposited by David Liu laboratory). Plasmids should be cited as gifts from David Liu with the corresponding Addgene plasmid number and RRID.

**Supplementary Table S6.**
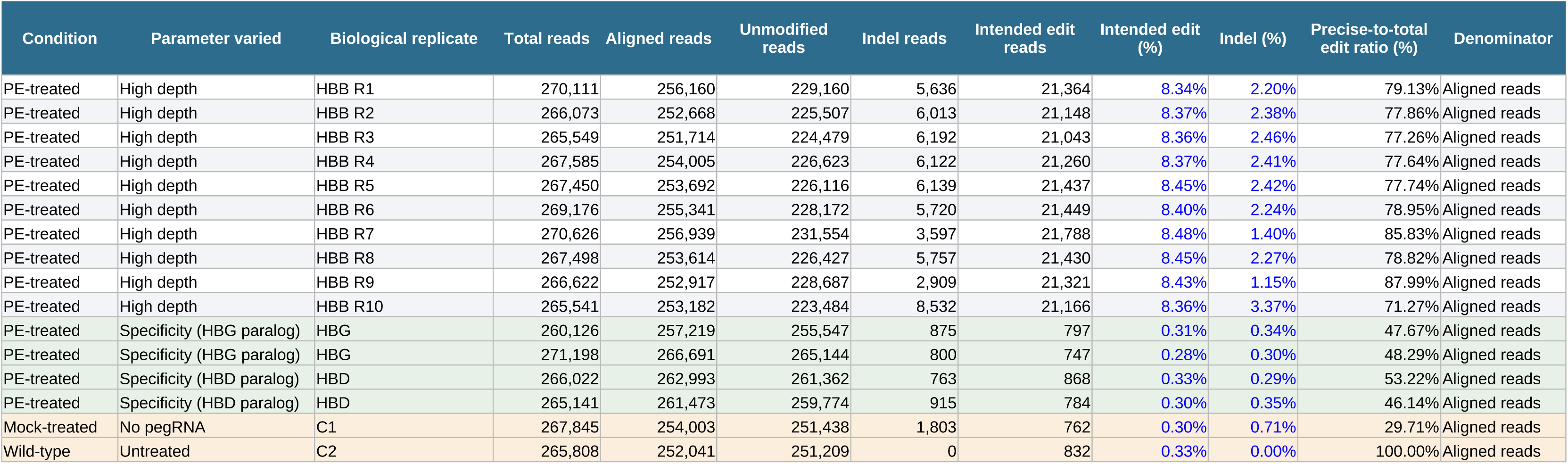
Prime editing outcome quantification at HBB codon 8/9 (c.27dupG)

## Notes

### Competing Interest Statement

The authors have declared no competing interest.

